# Expanding AHL Acylases Horizon - Insights From Structural Analysis of Choloylglycine Hydrolases From *Shewanella lohica* PV-4

**DOI:** 10.1101/788646

**Authors:** Pushparani D Philem, Yashpal Yadav, Avinash V Sunder, Deepanjan Ghosh, Asmita Prabhune, Sureshkumar Ramasamy

## Abstract

Acyl homoserine lactone acylases are quorum quenching enzymes that degrade the Gram negative bacterial autoinducer *N*-acyl homoserine lactone (AHL) and belong to the Ntn-hydrolases superfamily of enzymes. Recent findings reported AHL acylase activity of pencillin V acylases (PVA) which, alongside bile salt hydrolases, are members of the cholyolglycine hydrolase (CGH) family of the Ntn-hydrolases superfamily. The present study reports the unique activity profile of two CGHs from a marine bacterium *Shewanella loihica*-PV4, designated here as *Sl*CGH1 and *Sl*CGH2, including the structural analysis of *Sl*CGH1. Both the enzymes exhibit AHL acylase activity while unexpectedly being inactive on standard CGH substrates PenV and bile salts. *Sl*CGH1 differs from known homotetrameric CGHs in being a homodimer displaying a reduced active site volume attributed to loop orientation, which subsequently directs the substrate specificity. Moreover a ligand bound complex structure revealed an unusual bent conformation of the saturated acyl chain bound to the active site and also predicts a single oxyanion hole forming residue during catalysis instead of the usual two residues. Phylogenetic analysis reveals *Sl*CGH1 homologs cluster separate from reported CGHs and AHL acylases. On the whole, *Sl*CGH1 could represent a functionally distinct new sub-class of CGH as an adaptation to the marine environment and its structure could provide the structural framework for understanding such a novel subclass. We also make a modest proposal of a probable evolutionary link between AHL acylases and β lactam acylases based on the overlap in activity and structural features.

**Significance:** Cross-reactivity between AHL acylases and b lactam acylases has been recently identified giving us a vivid glimpse of a possible evolutionary relationship between the phenomena of quorum sensing and antibiotic resistance. We report here the first AHL acylase of the CGH structural framework. *Sl*CGH1 from *Shewanella loihica* PV-4 is also the first report of a marine CGH with a unique activity and a new structural subclass of CGH family with AHL acylase activity. This finding highlights the vast diversity of AHL acylases and by extension quorum quenching enzymes as adaptation to different habitats. The results from this study also bolster the link between signal molecules and antibiotics, extending our understanding of the inadequately understood physiological roles of b-lactam acylases.

---

Substrate promiscuity or cross-reactivity among different members of an enzyme superfamily can be useful tool to decipher functional and evolutionary relationships (1). The quorum quenching acylases acting on *N-*acylhomoserine lactones (AHLs) belong to the Ntn hydrolase superfamily, together withβ−lactam acylases such as penicillin G acylases (PGA), Penicillin V acylases (PVA) and cephalosporin acylases (CA). Recent findings (2–5) reveal structural similarity and a significant overlap in the substrate spectra of these enzymes, providing insights intoprobable links to bacterial signaling and antibiotic resistance (6).

AHLs, produced by Gram-negative bacteria, are one of the most extensively studied signal moleculesfor bacterial communication or quorum-sensing (QS, 7,8). AHLacylases (catalyse amide bond hydrolysis) and lactonases (catalyse lactone ring hydrolysis) are the two major classes of enzymes that degrade AHLs, thus involved in quorum quenching (9–11).While AHL lactonases belong to three different enzyme superfamilies (12), a majority of AHL acylases belong to the Ntn hydrolase superfamily.

Canonical AHL acylases, as Ntn hydrolases, are characterized by the presence of the αββα fold in their tertiary structure housing the N-terminal catalytic serine nucleophile, and cleave the amide bond between lactone ring and the fatty acyl side chain in AHLs (13). AHL acylases are closely related to PGA and CA structurally with a heterodimeric subunit composition and similar catalytic mechanism;substrate cross reactivity has also been reported between AHL acylases and PGAs. For instance, PGA from *Kluyvera citrophila* (*Kc*PGA, 4) can degrade C_6_-C_8_ HSLs and an AHL acylase produced by *Streptomyces* sp. (AhlM, 2) could degrade PenicillinG. Moreover, in a recent report, the bifunctional role of an acylase MacQ (from *Acidovorax* sp. MR-S7) has been suggested to mediate both quorum quenching and beta-lactam antibiotic resistance, owing its ability to degrade AHLs and β−lactam antibiotics (14).

On the other side, PVAs have also been recently shown to exhibit activity on long chain AHLs (5), although they are Cys-Ntn hydrolases and are sequence-wise distant from PGAs and canonical AHL acylases (15). PVAs are structurally and evolutionarily related to bile salt hydrolases (BSH), which are involved in the deconjugation of glyco-and tauro-conjugated bile salts in mammalian gut bacteria. Together PVA and BSH constitute the cholylglycine hydrolase (CGH) family. While the substrate set of CGHs has been so far limited to Penicillin V and bile salts, it can now be extended to include AHLs as well. All these findings reveal that acylase activity on AHLs might be spread across a wider spectrum of enzymes.

In our attempt to characterize a marine CGH homolog of the Ntn hydrolase superfamily, we encountered the existence of a unique activity profile, supplemented by structural variations, which could enhance our understanding of the evolutionary relationship between AHL acylases and β−lactam acylases. *Shewanella loihica*-PV4 is a strain isolated from iron-rich microbial mat of Loihi Seamount in the Pacific Ocean (16), and contains in its genome two putative CGH genes with low sequence homology (<25%) from known CGHs. Here, we report enzymatic activity of these two homologs, Shew_0681 (annotated as PVA-like enzyme) and Shew_2805 (annotated as cholylglycine hydrolase), designated henceforth as *Sl*CGH1 and *Sl*CGH2, respectively. Both enzymeshave atypical homodimeric structures and are specific for AHLs, while being inactive on β−lactam antibiotics and bile salts, thereby opening upthe spectrum of CGHs and AHL acylases that could exist in different habitats. Structural analysis of *Sl*CGH1and sequence clustering indicates that these acylases might have evolved as quorum quenching enzymes adapted to the marine environment.

## Results and Discussion

The marine psychrophilic bacterium *S. loihica* possesses two sequences in its genome with low homology to known CGH enzymes. *Sl*CGH1and *Sl*CGH2 exhibit 23% homology with the recently characterized PVA from the Gram-negative *Pectobacterium atrosepticum* (17), and 35% sequence similarity with each other. However, the residues deemed important for CGH catalytic mechanism (18) are strictly conserved. A BLAST search of both enzymes revealed multiple homologous sequences from marine bacteria, dominated by *Vibrio* and *Agarivorans* with few from *Shewanella*. The presence of two CGHs was observed in many members of the genus *Shewanella*. Given the evolutionary distance from other CGHs and the marine habitat, we reasoned that the *S. loihica* acylases could exhibit significantly different features from known CGHs.

Both *Sl*CGH1and *Sl*CGH2were expressed as soluble cytoplasmic proteins in *E. coli*. While *Sl*CGH1 showed high protein yield (200 – 250 mg/l), *Sl*CGH2 was rather poorly expressed (0.2 - 0.5 mg/l). Interestingly, both the acylases were completely inactive on penicillins (Pen V and G) and bile salts, which are canonical substrates for CGH. Besides, they also did not show any activity on β-lactam substrates including ampicillin, cephalosporin, carbenicillin as well as other amide bond substrates such as casein and p-nitrophenyl palmitate. Although inactive BSHs have been reported in *Lactobacillus plantarum* WCFS1 (19), these are often isoforms of a principal active BSH enzyme.Regardless, these observations reflect a gap in sequence based understanding of CGH function in general and a need to extensively explore functional perspectives of this enzyme family. Moreover, as revealed by gel filtration chromatography and MALDI-MS (Fig. S1A and S1B), *Sl*CGH1 also differs in oligomeric state from previously reported CGHs. It was expressed in solution as a homodimer, while known CGHs are usually homotetrameric in nature. Although PVAs from Gram-negative bacteria exhibit reduced oligomeric interactions, they still retain the tetramer arrangement (20), while *Sl*CGH1 is dimeric. These deviations in activity and quaternary structure hinted at the possibility of existence of a discrete group of CGHs, driving us to explore further.

### *Sl*CGHs show robust activity on long chain AHLs

In a recent report, PVAs from Gram-negative bacteria, *Pectobacterium atrosepticum*(*Pa*PVA) and *Agrobacterium tumefaciens* (*At*PVA) have been reported to also degrade long chain AHLs (5). Considering the sequence similarity, the AHL degradability of *Sl*CGH1and *Sl*CGH2was also assessed by studying the quenching of bioluminescence in biosensors, *E. coli* pSB401 and *E. coli* pSB1075, covering a wide spectrum of unsubstituted and 3-oxo-AHLs from C_6_ to C_12_-HSLs. *Sl*CGH1 exhibited maximum activity against 3-oxo-C_10_-HSL followed by C_10_-HSL (Table 1). At a lower concentration of AHLs (15 µM), *Sl*CGH1showed moderate activity against C_8_-HSL and 3-oxo-C_8_-HSL as well (data not shown). On the other hand, *Sl*CGH2 showed robust activity on AHLs with acyl chain longer than 10 carbons, with maximum activity on 3-oxo-C_14_-HSL, followed by C_14_-HSL. Mild activity was also observed on 3-oxo-C_8_-HSL, C_10_-HSL and 3-oxo-C_10_-HSL; both the enzymes did not act on C_6_-HSL (Table 1). Although *Sl*CGH activity was optimum against 3-oxo-HSLs, the preference for 3-oxo-substituted substrate did not form a consistent pattern with the rest of the AHLs tested. The nature of AHL degradation by *Sl*CGHs was further investigated using High Resolution Mass spectroscopy (HR-MS) to detect the catalysis reaction products. Mass spectra revealed peaks at *m/z* 124 and 102 corresponding to molecular formula of sodiated and protonated adducts of HSL, C_4_H_7_O_2_NNa and C_4_H_8_O_2_N respectively, demonstrating acylase activity (Fig. S2). An AHL acylase, Aac, from *Shewanella sp*. MIB015 isolated from fish intestine has been previously reported for its ability to degrade long chain AHL (21).

**Table 1.** Result of AHL degradation by *Sl*CGH1, C1S variant and *Sl*CGH2 assayed using Lux-based biosensors (*E. coli* pSB401 and *E. coli* pSB1075). Residual RLU % = RLU of test/ RLU of control*100 (mean± SD).

| AHLs | Residual RLU % |  |  |
| --- | --- | --- | --- |
|  | *S/CGH1* | C1S | *S/CGH2* |
| **C<sub>6</sub>-HSL** | 107.50 ± 14 | 129 ± 1 | 100.32 ± 4.3 |
| **C<sub>8</sub>-HSL** | 93.77 ± 10.15 | 130 ± 16 | 114.86 ± 9.8 |
| **3-oxo-C<sub>8</sub>-HSL** | 97.69 ± 9.8 | 92.3 ± 2.6 | 94.19 ± 3.9 |
| **C<sub>10</sub>-HSL** | 25.13 ± 6.8 | 89.3 ± 7 | 93.14 ± 2.8 |
| **3-oxo-C<sub>10</sub>-HSL** | 0.91 ± 0.2 | 89.1 ± 8 | 95.93 ± 2.3 |
| **C<sub>12</sub>-HSL** | 95.80 ± 2.3 | 85.2 ± 5 | 13.65 ± 6.8 |
| **3-oxo-C<sub>12</sub>-HSL** | 94.0 ± 5.5 | 84 ± 8.1 | 39.36 ± 4.9 |
| **C<sub>14</sub>-HSL** | 119.00 ± 11 | ND | 9.41 ± 3.003 |
| **3-oxo-C<sub>14</sub>-HSL** | 82.77 ± 0.83 | ND | 1.06 ± 0.67 |

These activity profiles of *Sl*CGH1 and *Sl*CGH2 are unique in that the enzymes are active solely on AHLs, although they show sequence similarity and conservation of catalytic residues corresponding to CGHs (Fig. 3E). With respect to the AHL specificity, a preference for long chain AHLs (C_8_ to C_14_-HSL) is common among different acylases, including PVAs. Heterodimeric AHL acylases, PvdQ and QuiP from *P. aeruginosa* PAO1 (22, 23), AhlM from *Streptomyces* sp. strain M664 (2), AiiC from *Anabaena* sp. PCC7120 (24) and QqaR from *Deinococcus radiodurans* R1 (25) showed preference for AHLs with acyl chain longer than 6 carbons. *Pa*PVAwas strictly active on C_10_ and C_12_-HSL (5), while *At*PVA was active on a wider range of AHLs with maximum activity on long chain AHLs. Unlike these examples, AiiD from *Ralstonia* sp. JX12B (26) can hydrolyze C_4_-HSL as efficiently as it degrades 3-oxo-C_12_-HSL, while it shows significantly low activity on 3-oxo-C_6_-HSL. MacQ from *Acidovorax* sp. MR-S7 (14) displays a broad spectrum of activity, degrading C6 to 3-oxo-C_14_ HSL. PGA from *Kluyvera citrophila* is most active on 3-oxo-C_6_ HSL (4). Multiple AHL acylases produced by a single species can also exhibit differential substrate specificity as in the case of *S. loihica*;this is exemplified byHacA (C_8_-C_12_ HSLs) and HacB (C_6_-C_12_ HSLs)from *Pseudomonas syringae* (27).

**Fig. 3.**
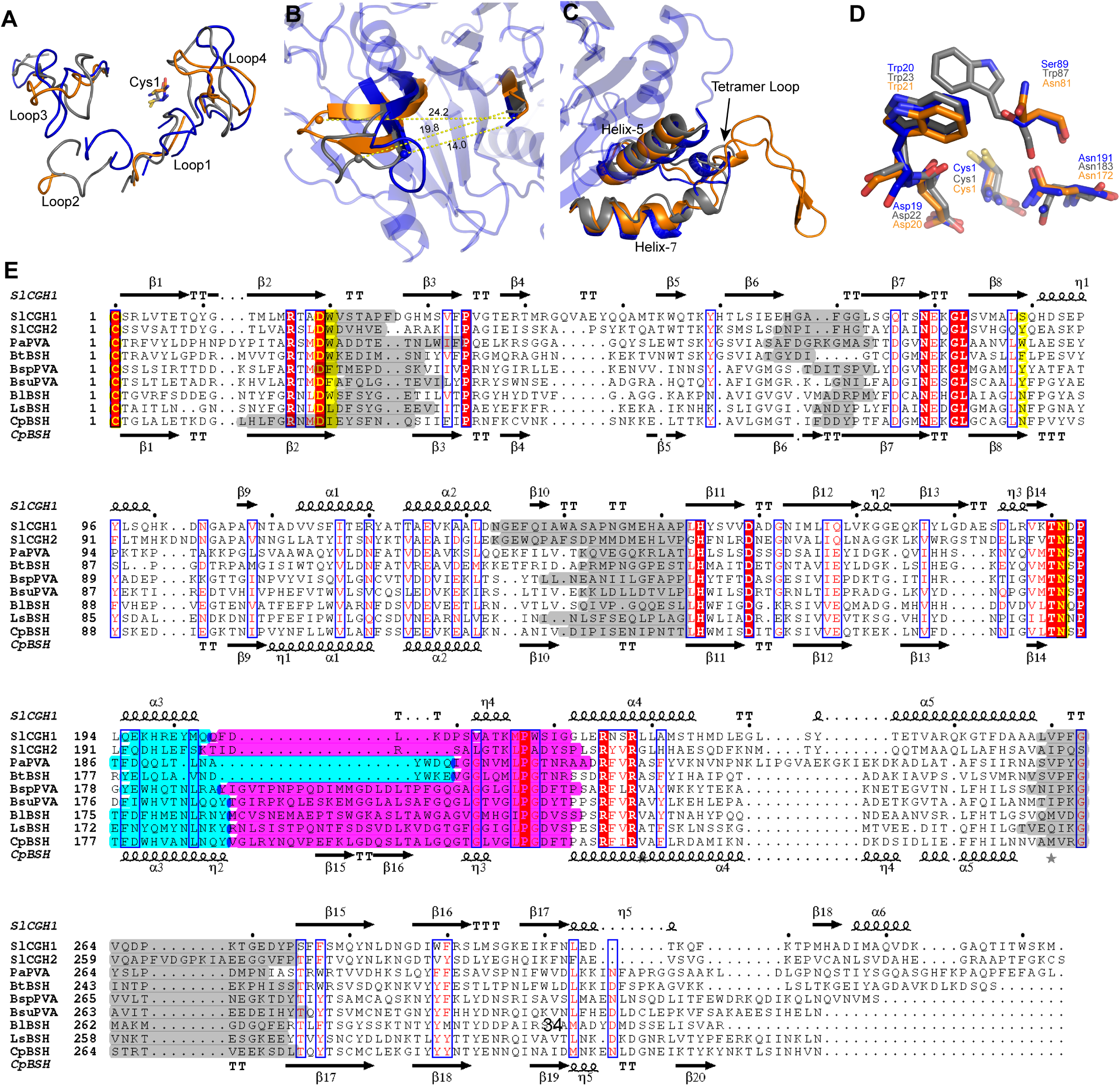
Similarity and differences in CGH family *Sl*CGH1 (blue), *Pa*PVA (gray) and *Bl*BSH (orange). A) Superposed conserved active site loops (Loop1 to Loop4) in *Sl*CGH1, *Pa*PVA and *Bl*BSH highlighting difference in orientation of loop2. B) Distance between centroid of the loop2 and Cys1, showing inward orientation of *Sl*CGH1 loop2. C) A short tetramer motif/loop is flanking between α5 and α7 in Gram negative CGH (*Sl*CGH1and *Pa*PVA) as compared to *Bl*BSH. D) Superposed conserved active site residues represented with stick. Alignment coloring are based on ESPript (http://espript.ibcp.fr) While most residues remain conserved, one of the oxyanion forming residues varies from Trp87in PVAs, Asn81 in BSHs to Ser89 in *Sl*CGH1. E) Sequence alignment of CGH enzymes. *Sl*CGH1, *Sl*CGH2, PVAs from *Pectobacterium atrosepticum* (PaPVA), *Bacillus subtilis* (*Bsu*PVA) and *B. sphaericus* (*Bsp*PVA); BSHs from *Bacteroides thetaiotamicron* (*Bt*BSH), *Bifidobacterium longum* (*Bl*BSH) and *Lactobacilus salivarus* (*Lb*BSH) and *Clostridium perfringens* (*Cp*BSH). The alignment highlights conserved residues (red), loops (grey) flanking the active site (yellow) and assembly motif (Magenta), and α helix involved in oligomeric interactions between two monomeric subunits (cyan).

### Quaternary structure of *Sl*CGH1 reveals an atypical dimer

The apo-*Sl*CGH1 crystal structure, solved at 1.8 Å resolution using X-ray crystallography, reveals the presence of the characteristic αββα-fold of Ntn-hydrolase superfamily (Fig.1A). The enzyme crystallized in the I12_1_ space group with two *Sl*CGH1molecules per asymmetric unit (Table S2 and S3). In the case of *Sl*CGH2, thin needle crystals were obtained in the Index screen (Hampton,USA). However, the crystals failed to diffract and crystal quality could not be improved despite several rounds of manual optimization. As a result, further structural analysis was restricted to *Sl*CGH1.

**Fig. 1.**
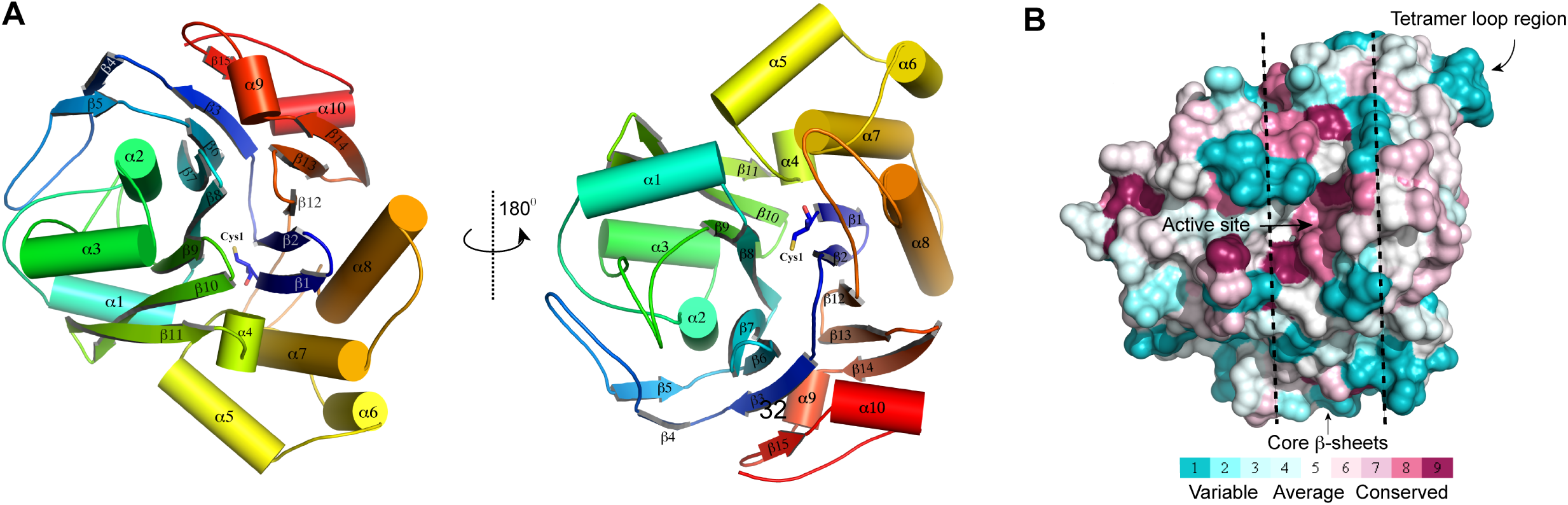
Crystal structure of *Sl*CGH1. A) Cartoon representation of secondary structural elements in *Sl*CGH1 monomer colour-ramped from N- to C-terminus (blue to red). The α-helices and the β-strands are labelled and rotated at 180° B) ConSurf analysis (55) of *Sl*CGH1. Molecular surface representations are colored by their conservation score higly conserved regions (maroon) and variable regions (cyan) and dotted lines indicate the core β-sheets.

**Table 2.**
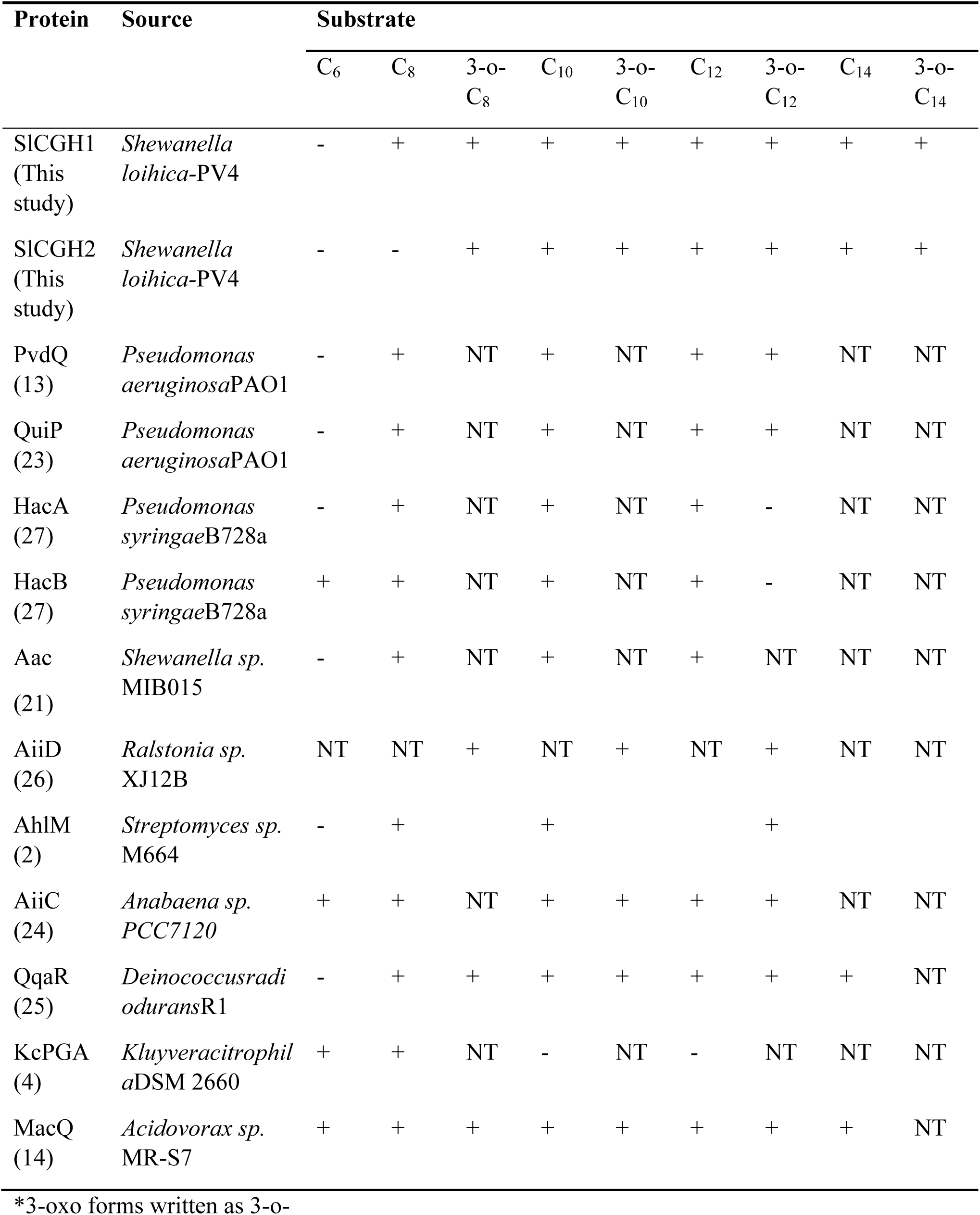
List of known AHL acylases and their activity profile.

In Ntn hydrolases, two core β sheets host the active site and are spatially conserved.The angle between these two sheets in *Sl*CGH1 is ∼23.78°, while previously reported CGHs show ∼30° separation(28). The consurf analysis shows conservation of active site and surrounding β-sheet while tetrameric loop region exhibits high variation (Fig. 1B). *Sl*CGH1 exhibits maximum structural similarity (DALI server) with CGHs from Gram negative bacteria, *Bacteroides thetaiotamicron* (*Bt*BSH) and *P. atrosepticum* (*Pa*PVA), followed by Gram positive CGHs (Table S4). The AHL acylase, PvdQ from *P. aeruginosa* shows a low identity of 7%, indicating a distant relationship between known AHL acylase and *Sl*CGH1.

*Sl*CGH1adopts a stable dimeric structure in both crystal form and in solution (Fig. 2A). PISA analysis of protein interfaces suggested that the dimer is thermodynamically stable, with a solvent accessible surface area of 25577.2 Å^2^ and buried surface area 2719.7Å^2^. The dimer interface in *Sl*CGH1isformed by the same regions (α6 and α8) of the Ntn fold as it occurs in other CGHs.Gram-negative bacterial CGHs have shown reduced dimer interface than their Gram-positive counterparts (6, 18); in *Sl*CGH1, the interface is spread across an area of 1359.8 Å^2^,which is the smallest value reported so far and is composed of mostly hydrophobic residues at one side and dominated by positively charged residues on the other (Fig. 2B). The two monomers are held by two hydrogen bonds along with 104 non-bonded contacts between 24 residues from both subunits (Fig. S3).

**Fig. 2.**
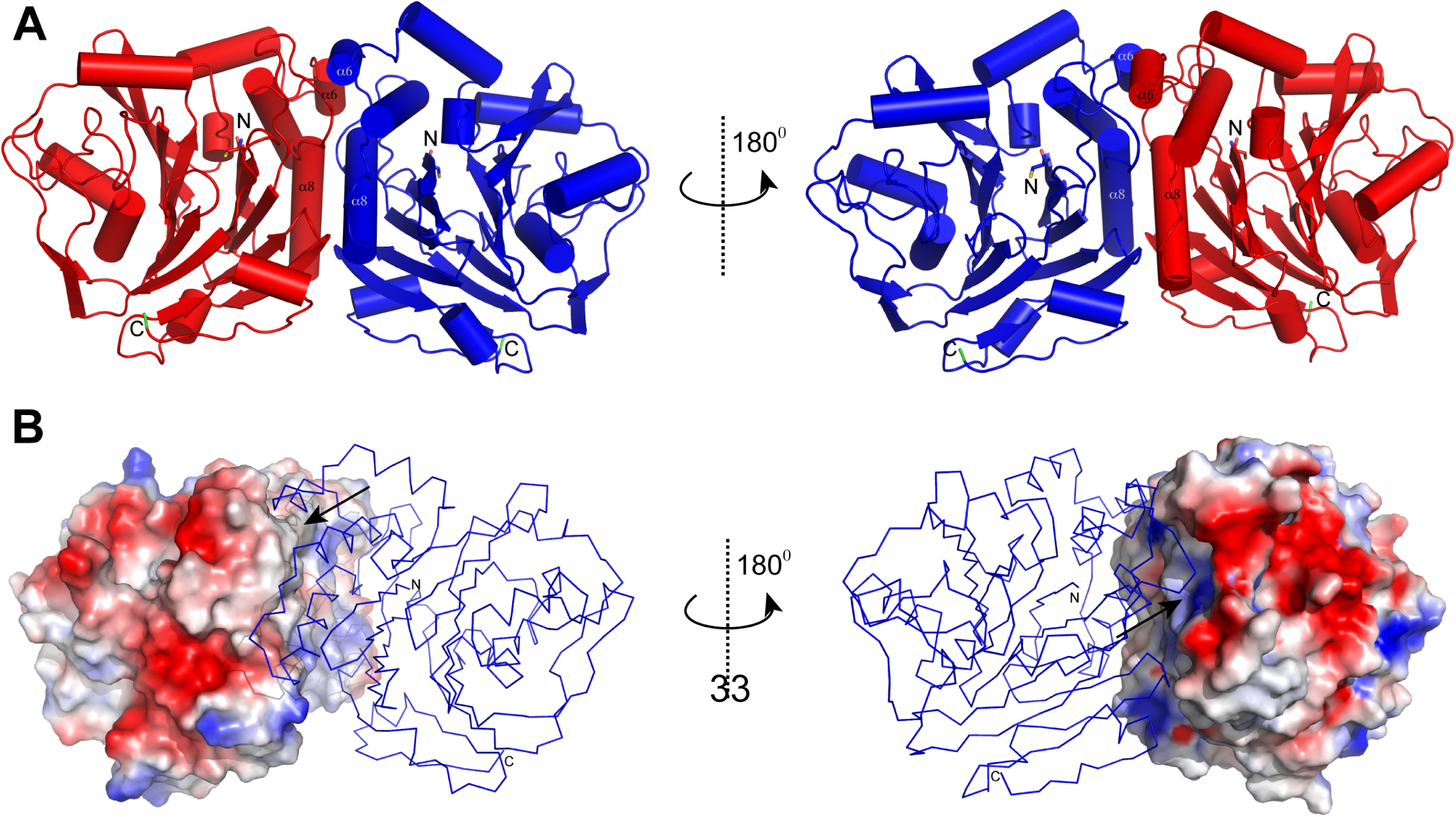
A) Overall structure of *Sl*CGH1 dimer shown in front view (left panel) with respect to Cys1 and rotated at 180° (right panel). B) Electrostatic surface potential of *Sl*CGH1 from negative (red) to positive (blue) and rotated at 180°. The arrow indicates the dimer interface is consist mostly hydrophobic (white) at one side and dominated by positive charge (blue) at other side.

A combination of features could contribute to the absence of tetramerisation in *Sl*CGH1. *Sl*CGH1 displays a short tetrameric loop, typical of Gram negative CGH, containing a small α helix (V212-K215) in the middle which might reduce its flexibility (Fig. 3C). Two α-helices on either side of the tetramerization loop in CGHs, corresponding to α5 (Q195-Q204) and α7 (G220-T234) in *Sl*CGH1, are important for interactions between the adjacent subunits of two dimers. In *Sl*CGH1, the exposed α5 helix consists entirely of polar residues, as opposed to a more even distribution of polar and non-polar residues in other CGHs. The change in polarity may increase the feasibility of α5 to remain exposed to solvents, thereby lessening the propensity to interact with another dimer for protection of non-polar residues. In addition, a shortened α5 (by four residues) in *Sl*CGH1distances potential interacting residues for a tetrameric alignment. In *Pa*PVA, two polar contacts (Y196:D214 and Q199:Q189) between these α-helices from adjacent subunits of the tetramer and a non-polar contact involving V204 and M205 from the tetramerization loops of opposite subunits constitute the main oligomeric interactions (20). Similar interactions were absent in *Sl*CGH1, as these interacting residues were either not conserved (Met) or positionally different (Val). We suggest that the difference in oligomeric state could reflect different evolutionary stages of CGH for functional diversification driven by adaptation to different environments.

### Loop conformational changes could explain *Sl*CGH1 activity profile

The active site of *Sl*CGH1 was considered from two perspectives – the catalytic residues and the loops surrounding the site. The structure was compared with CGHs from both Gram-positive and Gram-negative bacteria: *Pa*PVA (20), *Bsp*PVA (*Bacillus sphaericus*, 15), *Bt*BSH (*Bacteriodes thetaiotamicron*, 29) and *Bl*BSH (*Bifidobacterium longum*, 28). The *Sl*CGH1active site shares multiple conserved residueswithknown CGHs such as R16, D19, W20, N191 and R225 involved in catalytic mechanism andsubstrate binding (Fig. 3D). The only exception concerned the oxyanion forming residue S89 which corresponds to Tyr or Trp in PVAs (15, 20) and Asn in BSH (30). However, this variation might not impact the nature of catalysis, as it is the backbone amide group that participates as electron acceptor during the formation of the intermediate oxyanion hole (30). Other homologous acylases of marine origin, including *Sl*CGH2 possess a Tyr or Trp at this site.The active site is predominantly hydrophobic due to the flanking aromatic residues W20, F26, F70 and F276 from the surrounding loops and hydrophobic residues such as A18, I64, A87, L88, L153 etc. from the surrounding β sheets.

In CGHs, four conserved loops flanking the active site (Fig. 3A) are important for substrate binding, with the differences in loop length and conformation affecting the size and properties of the active site and substrate binding pockets (28, 18). Although the core active site residues remain largely conserved in *Sl*CGH1, the active site loops show significant deviations in position and length compared to other CGHs. Loop 2 in particular displays an inward shift, moving closer to Cys1 by ∼5 Å and 9 Å when compared to *Pa*PVA and *Bl*BSH, respectively (Fig. 3B). Consequently, the active site shrinks to a much smaller volume (79 Å^3^) than in any CGH reported so far (153-718 Å^3^, 18). It is possible that this might have been facilitated by the presence of four Gly residues (G68, G71, G72 and G75) in *Sl*CGH1 loop2. Besides reducing the active site volume, this also creates a narrow, close ended U-shaped architecture of the binding pocket as loop 2 and 3 move closer on one end of the pocket. We suggest that this narrow active site might allow entry of substrates with a linear geometry such as fatty acid chain in AHL, while blocking entry of bulky substrates such as Pen V (phenyl ring) and bile salts (steroid moiety) and hence the inactivity towards these substrates.

In addition, loop3 in *Sl*CGH1was extended by 4-14 residues than in previously characterized CGHs (Fig. 3E), primarily due to the disappearance of a short β-strand present in other CGHs. It also exhibits high average B factor with exceptionally highest value (31.2 Å^2^) centered on residues 147-149, which further increases in the ligand-bound structure (40.4 Å^2^) (Fig. S4). As B-factor represents real molecular motions, these higher values support the extra flexibility of loop3 as against the rest of the protein which mightindicate the possible function of loop3 residues in ligand entry or exit.

### Structure of C1S:3-oxo-octanoic acid reveals an unusual bent acyl chain conformation

Attempts to crystallize the enzyme-substrate complex using both native *Sl*CGH1and site-directed (Cys1) mutants yielded a C1S complex with bound 3-oxo-octanoic acid (3OOA) product. The 3OOA chain was trapped in the groove between β sheets I and II flanked by loops1-3 (Fig. 4A), corresponding to the general Pen V /bile salt binding site in CGHs. The acyl chain was bent in the middle between C3 and C4 atoms, probably due to obstruction by β5 and β6 strands (Fig. 4B). This is rather unusual as bent acyl chain conformation is generally associated with flexible unsaturated acyl chains, where it improvestight substrate binding and catalytic efficiency (31). Unlike *Sl*CGH1,PvdQ accommodates an elongated acyl chain conformation in a deep hydrophobic binding pocket (13). Strikingly, the distances between carboxylate oxygen of the 3-oxo-octanoic acid and the backbone amide of S89 and Nδ of N191 in the *Sl*CGH1 complex structure are 2.8 and 4.1Å, respectively, which is almost a precise match with the corresponding values in PvdQ, 2.8/4.2Å (Fig. 4B).

**Fig. 4.**
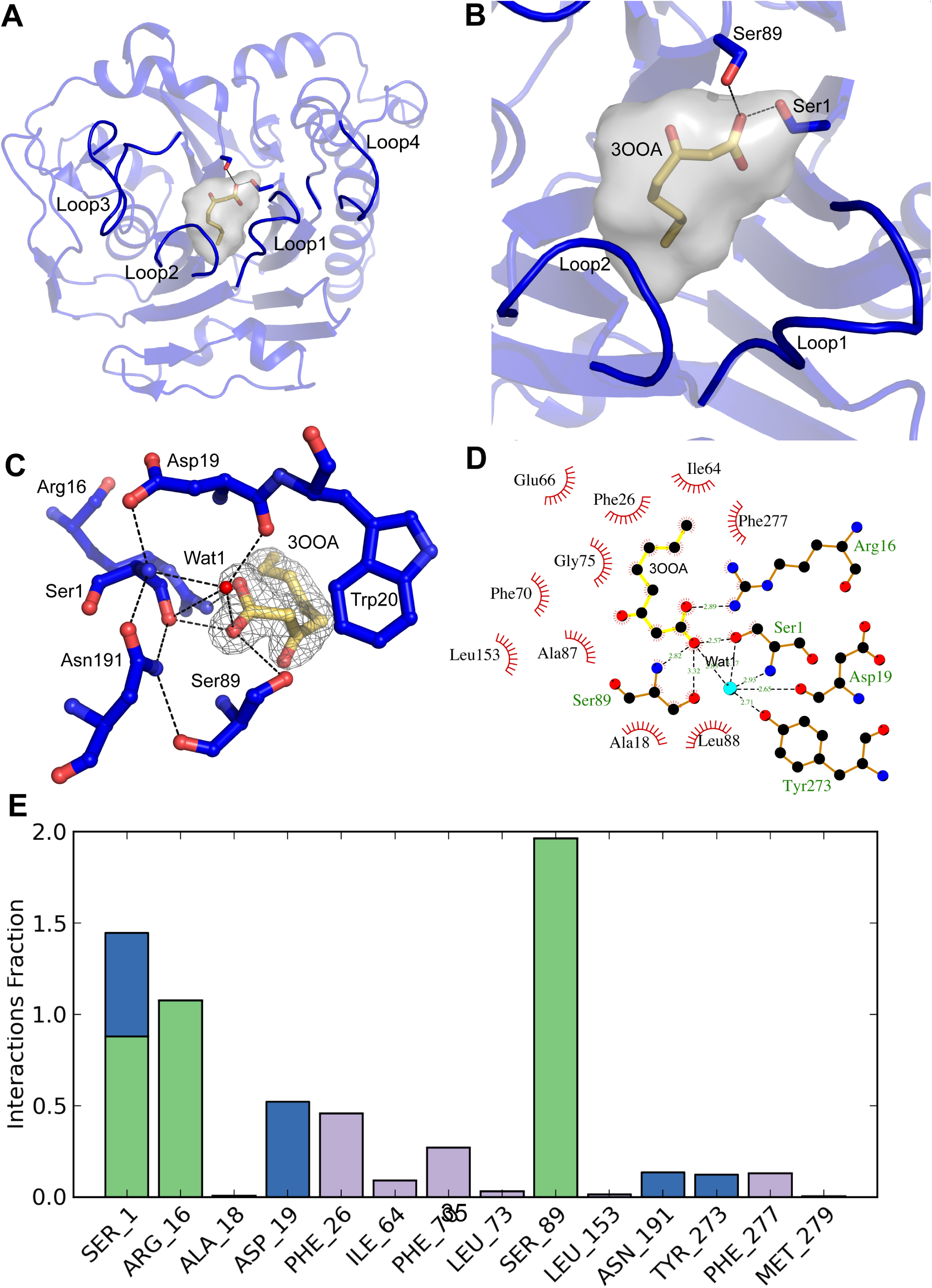
Co-crystal structure of *Sl*CGH1 with 3OOA A) Structure of C1S-OOA complex showed the hydrolysed product, 3OOA, bound in the hydrophobic groove flanking the active site loops. B) Zoomed in view of binding site of 3OOA with interactions also shows an unusual bent conformation of the acyl chain between C3 and C4 atoms. C) Electron density contoured (gray mesh) around 3OOA at 1σ. D) Ligplot scheme showing interactions within the complex. H-bonds are presented as black dashed lines with corresponding length. 3OOA makes strong H-bonding with Ser89 at the two positions. The water molecule (cyan) near Cys1 makes an extensive of H-bonding network. e) Histogram showing protein-ligand interactions from MD simulation of C1S-OOA complex structure. The interaction is dominated by H-bond (Green bar) involving Ser1, Arg16 and Ser89, followed by mild hydrophobic bond (Purple bar) and water bridges (Blue bar). No H-bonding is contributed by Asn191 which is the expected oxyanion residue.

Superposition of the ligand-bound C1Svariant on apo-*Sl*CGH1structure reveals a low RMSD of 0.083 Å (Fig. S5). There is a slight increase in the whole chain average B-factor of the complex structure (20.09 Å^2^) than that of the apo form (14.96 Å^2^). It could therefore be assumed that the AHL binding in *Sl*CGH1follows mild conformational changes in the enzyme structure. However, it is also possible that since only the hydrolyzed product was found bound to the enzyme, the complex structure could represent a captured snapshot after the catalytic reaction when the enzyme has returned to its native form. The complex structure further reveals that the hydrolyzed product (3OOA) was bound to the active site with one of its carbonyl oxygen in polar contact with a water molecule and the OH group of Ser1 (Fig. 4C). At this stage the amino part of the substrate, here the lactone ring, has left the active site. This depicts the second tetrahedral intermediate, formed after the cleavage of amide bond, wherein the water molecule acts as the virtual base to cause the second and final nucleophilic attack to release the acyl group (32, 33). Moreover, no specific interactions were observed between the 3-oxo group of the AHL substrate and the enzyme, which is in agreement with the report on PvdQ (13). It has been suggested that the polar oxo group might ease the exit of the acyl chain out of the hydrophobic binding pocket (13). Structural comparison between *Sl*CGH1and the AHL acylase PvdQ (13) was difficult as the two enzymes are oligomerically different, despite similar activity profiles. The active site of *Sl*CGH1(79 Å^3^) is smaller than that of PvdQ (144 Å^3^). However, the core β sheets I and II that host the active site are highly conserved structurally. Superposition of only these β sheets revealed the conservation of core active site architecture, albeit with different residues (Fig. 5A).The oxyanion hole forming residues in PvdQ, V70 and N269 correspond to S89 and N191 in *Sl*CGH1. CASTp analysis shows an open active site groove while *Sl*CGH1 exhibits the closed one (Fig. S6).

**Fig. 5.**
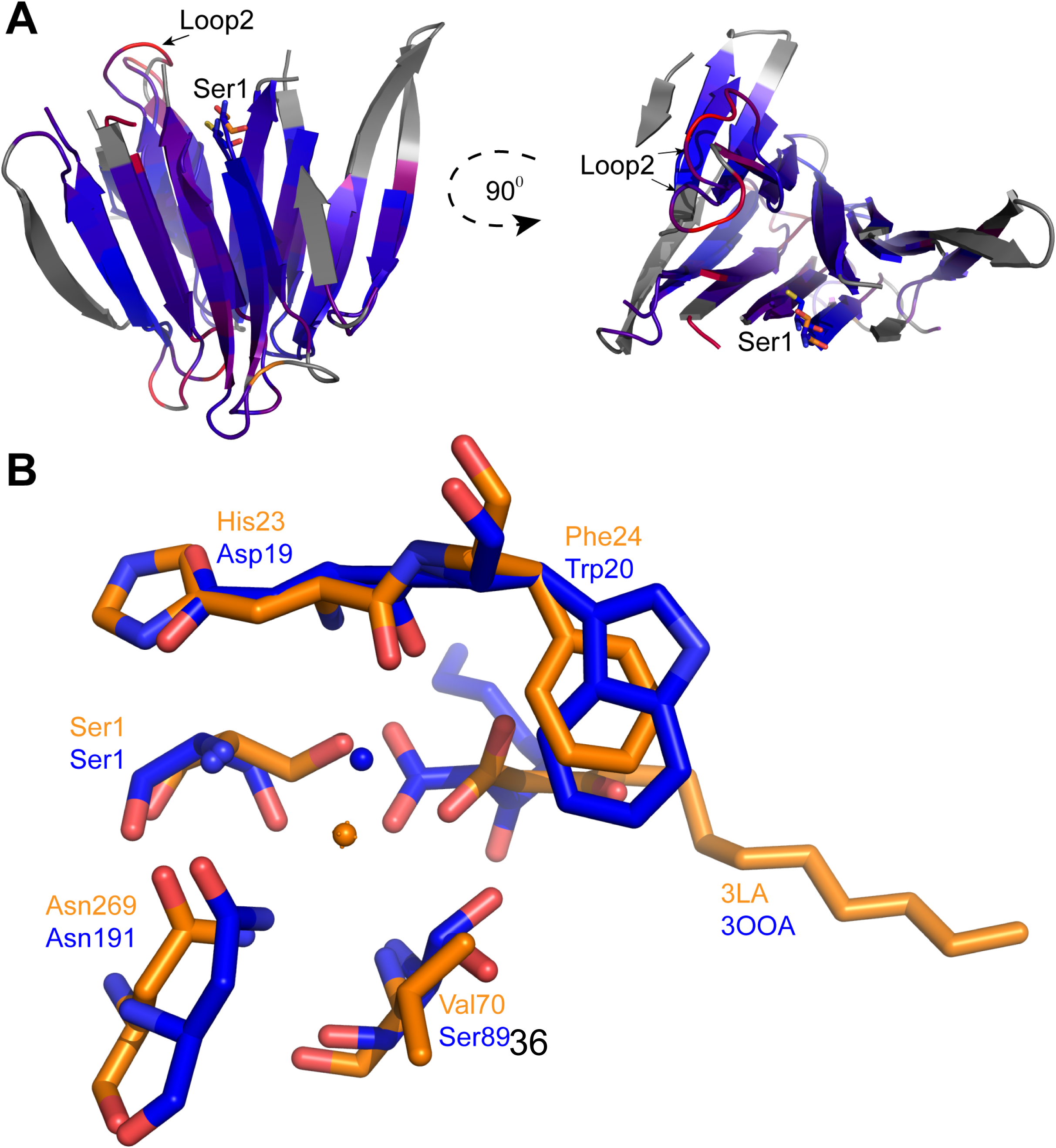
Comparision between crystal structures of *Sl*CGH1 and PvdQ A) Conservation of core β-sheets (front and rotated at 90°) between *Sl*CGH1 and PvdQ is illustrated by superimposing the same regions and colored by RMSD with color ramped from blue to red with increase in RMSD. Regions in grey are not considered for superposition. B) Superposed view of ligand bound active sites – C1S (blue) shown with 3OOA and PvdQ (orange) with 3LA. The conserved water molecules near Ser1 and the corresponding conserved residues were indicated.

In the ligand bound C1S variant, the carbonyl oxygen atom of the acyl chain is hydrogen bonded with S89 at two positions, hydrogen of the backbone amide and the hydrogen of the hydroxyl side chain (Fig. 4D). In MD simulation study, hydrogen bonding interactions were observed throughout the simulation time scale (1.2 ns) between the residues S1, R16, S89 and the ligand OOA in *Sl*CGH1. S89 forms hydrogen bond with the ligand throughout the entire simulation time scale while N191 shows no hydrogen bonding with the ligand except weak intermittent water bridges (Fig 4E). This observation raises the possibility of S89 alone participating in the oxyanion hole formation for the stabilisation of the tetrahedral intermediate, instead of the predicted pair of S89-N191. N191 rather interacts with S1and S89, stabilizing their negative charges. MD Simulation study further reveals a solvent accessible Surface Area (SASA) for the ligand to be 15-fold lesser than that of 3-oxo-dodecanoic acid bound PvdQ, highlighting the significant difference in the overall nature of the active sites in the two enzymes. This observation is also very interesting given the highly hydrophobic active site of PvdQ (13).

The N-terminal Cys1 as the catalytic nucleophile in *Sl*CGH1was validated from the loss of AHL acylase activity in the C1A mutant. However the C1S variant retained mild activity on long chain AHLs (Table 1), indicating a similar mechanism of catalysis between Ser- and Cys-AHL acylases. Conversely, in a previous report on PGA from *E coli* ATCC11105, a mutation of nucleophile Ser to Cys resulted in loss of activity (34) whichpoints towards a more distant structural relationship between PGA and PVA. The catalytic mechanism in Cys and Ser Ntn-hydrolases, in general, has been reported to show subtle differences, mainly due to the nucleophile differences while the overall catalysis mechanism stays the same (15). In *Sl*CGH1, a water molecule is observed in both the apo and complex structure, at hydrogen bonding distance from both the Oγ (3.2 Å) and the α-amino group (2.9 Å) of the nucleophile. Owing to its almost neutral side chain pKa, Cys does not require a catalytic base like water for nucleophile activation as required by Ser-Ntn-hydrolases, (15, 35, 36), even though a water molecule is found conserved in both the type of enzymes (Fig. 5B).The reaction starts with a direct proton transfer from the Sγ to the α-amino of Cys1 leading to the activation of the nucleophile (Fig. S7).

Docking of C_6_-HSL and C_8_-HSL showed lower Glidescores than that of 3-oxo-C_10_-HSL. The distance of the carbonyl carbon of the amide bond from the SH of the Cys nucleophile (nucleophilic attack distance, NAdist) is also favourably the lowest for 3-oxo-C_10_-HSL (Table S5, Fig. S8). These factors could explain the high activity of *Sl*CGH1on 3-oxo-C_10_-HSL, as indicated by the bioluminescence assay (Table 1). Although Glide scores were similar for C_6_-HSL and C_8_-HSL, the farther NAdist for C_6_-HSL might lead to unproductive binding, whereas moderate activity could be detected on C_8_-HSL at lower AHL concentration. C_8_-HSL and C_6_-HSL were observed to bind in a similar orientation in the active site making a bend at the C3 of the chain. Inside the active site cavity, shorter acyl chain ends in a surrounding dominated by hydrophilic residues, E65, E66, H67 and Q76, which may lead to weaker binding. Longer acyl chains can access the more hydrophobic interior part of the pocket, lined primarily by residues I64, F70, L73 and M279 (Fig. S8b). It can be inferred that the presence of extended acyl chain might lead to increased hydrophobic interactions in the binding pocket, thus allowing productive binding and better AHL degradation activity, similar to *Pa*PVA (5). With 3-oxo-C_10_-HSL, there is a sharp bend at C3 atom of the acyl chain orienting the lactone ring closer to the Cys nucleophile in *Sl*CGH1. The docked AHL ligand binds in a similar orientation as the acyl chain in the C1S:3OOA complex structure, although the acyl chain in the complex structure was captured closer to the nucleophile (3.3 Å away from Ser1). There were no polar contacts between the lactone ring of the AHL substrate and the active site residues.

### *Sl*CGH1shows phylogenetic and structural differences from other CGHs

The phylogenetic separation of CGHs of Gram positive and Gram negative origin in two different clusters has been analyzed recently (18). Extending the analysis using homologs obtained from BLAST analysis of *Sl*CGH1 revealed theirsegregation into a distinct third clade/cluster (Fig. S9A). However, *Sl*CGH1 hits were scarce (82) and almost all homologs were sequences from marine dwelling bacteria. Moreover, it was also surprising that there were only 3 sequences from *Shewanella* sp. in the list, including *Sl*CGH2. The first two best hits (identity score of 72-66 %) were un-annotated hypothetical proteins from *Vibrio fluvialis* and *Agarivoran gilvus* respectively, while other hits had <40% identity. A large number of distant marine homologs could be identified with a relaxed E-score cutoff; however, these formed multiple smaller clusters on their own and were not included in the final phylogenetic tree to avoid complexity. These findings might indicate an adaptation to spatially different niches within the marine environment. Regardless, the phylogenetic information coupled with the unique specificity of *Shewanella* CGHs for AHLs (while inactive on Pen V or bile salts) provides intriguing information regarding the probable evolution of a new sub-class of CGHs in the marine habitat. This becomes more evident through the differences in subunit composition of the *Shewanella* CGHs and the structural features in the active site/substrate binding pocket which is more suited for AHL binding.

Panigrahi et al. (18) also employed the Binding site similarity (BSS) score matrix as an improved method of differentiation between PVA and BSH, based on the conservation of residues critical for PVA/BSH activity and substrate binding residues in the loops 1-4 surrounding the active site. A strict conservation of these residues with any of the six functional PVA/BSH templates would essentially translate to a higher BSS score, thereby providing conclusive information to categorize the CGH sequence as either a PVA or a BSH. However, on employing this algorithm with both *S. loihica* CGHs, no significant similarity could be achieved with any of the six templates used. This observation alludes to the fact that the CGHs of marine origin, including those from *S. loihica*, show significantly lower sequence homology with known CGHs from other environments and cluster separately. On adding *Sl*CGH1 as a new template to revise the BSS scoring system, high scores were obtained for some of the marine homologs with *Sl*CGH1. A few sequences (from *Photobacterium swingsii, Vibrio tubiashii,Shewanella waksmanii* and *Vibriomaritimus*) were ambiguous, showing BSS scores matching with all Gram-negative templates –*Pa*PVA, *Bt*BSH and *Sl*CGH1(Fig. S9B). It is possible that these enzymes might show some activity also with Pen V or bile salts.

### Diversity and evolution of acylases active on AHLs

*Sl*CGH1and *Sl*CGH2 add to the diversity of acylases that are now known to deacylate AHL signals. Their uniqueness stems from the fact that both are solely active on AHLs, lacking any activity on canonical CGH substrates Pen V or bile salts. Furthermore, this is the first report on CGH from marine bacteria. Marine CGH homologs were also observed to be evolutionarily distant from CGHs from both Gram-positive and Gram-negative bacteria, as evidenced by the formation of separate clusters. These findings suggest that CGH homologs from marine bacteria could represent a new sub-class of CGHs with unique substrate preference. CGHs so far have been reported to have substrate preference specific for Pen V (*P. atrosepticum*, *B. subtilis*) or bile salts (*Clostridium perfringens*), or show moderate cross-reactivity (*B. sphaericus* PVA, *Bifidobacterium longum* BSH) (28, 20). Recently, functional PVAs, highly active on Pen V, have been shown to also degrade long chain AHLs (5). Considering the activity profiles of *S. loihica* acylases and *Pa*PVA, we suggest that AHLs can also be added to the list of potential substrates for CGH enzymes.It should be noted though, that when we evaluated the AHL degradability of some BSHs, the BSH from *Enterococcus faecalis* (37) and a few other gut microbiota showed only low level activity on AHLs (Data not shown). The structural features of the BSH active site, including a hydrophilic loop3 and the presence of polar residues that form hydrogen bonds with the hydroxy groups of the steroid moiety in bile salts (38), could make the substrate binding pocket less hydrophobic and discourage the productive binding of AHLs. *L. plantarum* produces four BSH homologs with feeble activity on 3-oxo-C6 and 3-oxo-C8 HSLs (19); however, the inactive BSHs (bsh2-4) exhibited twice the activity on penicillins and AHLs than the principal bile acid-active enzyme (bsh1). Moreover, despite the complexity of bacterial communities in the mammalian intestine, the presence of AHL signalling among the normal gut microbiota has not been detected yet (39). It is therefore possible that the BSH might have evolved to shift the substrate specificity from AHLs to bile salts in the intestinal environment. Moreover, it might also be possible that such feeble activity is sufficient for modulating QS in the gut environment. In another scenario, the host bacterium might possess other enzymes with AHL degrading activity, besides BSH.

In the last decade, acylases that act on AHLs and mediate quorum quenching have been characterized from several bacteria. It is now clear that these acylases vary widely in their substrate spectrum and sequence, and separate into multiple phylogenetic clades (40, 14). Certain AHL acylases including PvdQ prefer long chain AHLs, while members of the QuiP group can act on both long and short chain AHLs, and the β-lactam acylases primarily act on Pen G. Some acylases are promiscuous, and can cleave both AHLs and Pen G. Such enzymes including AhlM and KcPGA are distributed across the branches of phylogeny. Another interesting case is the PVA from *Streptomyces lavendulae* (*Sl*PVA), which is a heterodimeric Ser-Ntn hydrolase that shares a significant similarity with AHL acylases and cleaves AHL; however, it was originally characterized as active on Pen V and penicillins with aliphatic side chains, with minimal activity on Pen G. Bacterial PVAs and the *S. loihica* acylases characterized in this study (Cys-Ntn hydrolases) that belong to the CGH group and can cleave AHL occur in a completely different phylogenetic branch, which adds another layer of complexity to the diversity of acylases acting on AHL signals (Fig. 6). It is important to note that some acylases were initially characterized on their ability to act on other substrates such as β-lactam antibiotics and bile salts, and data on their AHL degradation ability is often scarce. Comprehensive substrate profiles might help uncover the full functional range of these enzymes.

**Fig. 6.**
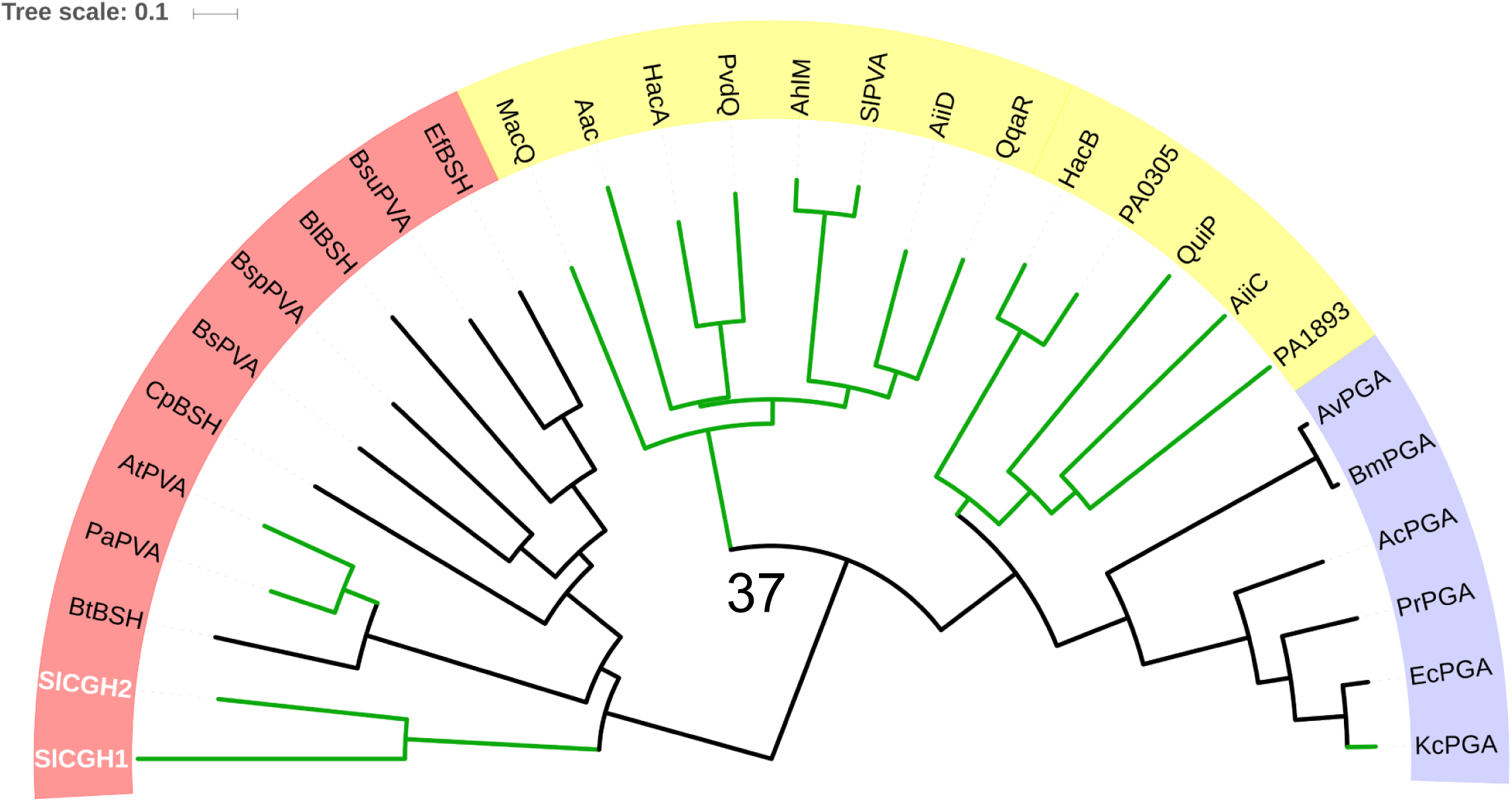
Phylogenetic anaylsis of different members of AHL acylases, PGA and CGH based on sequence similarity. All CGHs (red) forms a separate cluster from AHL acylase (yellow) and PGAs (purple). Green colored nodes indicate enzymes with positive AHL acylase activity. *Sl*CGH1 and *Sl*CGH2 form another separate clade among CGHs, close to AHL degrading CGHs of *Pa*PVA and AtPVA.

The diversity of AHL-active acylases and their related promiscuous substrates could be addressed from structural and evolutionary perspectives. While the core catalytic residues and the catalytic mechanism of Ntn hydrolases are strictly conserved, the shape and nature of the substrate binding pocket can often change,with subtle structural deviations altering the substrate spectrum. For instance, the PGA from *E. coli* lacks activity on AHLs in spite of 80% overall sequence similarity with *Kc*PGA, with the capacity of *Kc*PGA to productively bind AHLsattributed to the conformational variations in the active site residues β145R and β146Fduring enzyme processing (37).The specificity of PvdQ was shifted from long chain to medium chain AHLs through the mutagenesis of two residues (Lα146W, Fβ24Y) that decreased the size of the hydrophobic binding pocket (41). In the present study, we observed that changes in the residues and orientations of the loops flanking the CGH active site could modulate the substrate specificity between Pen V, bile salts and AHLs in different CGHs including *Sl*CGH1 and *Sl*CGH2.This serves to illustrate that the Ntn hydrolase fold allows for excellent versatility, allowing the enzymes to evolve through minimal structural modifications with changes in critical residues involved in substrate binding, resulting in a wide substrate spectrum.It has also recently emerged that enzymes possessing different folds might be AHL acylases too, such as α/β hydrolase (AiiO from *Ochrobactrum* sp., 42) or amidase (AmiE, 40).

Although a definite order of evolution for AHL-active acylases is not clear, the overlapping substrate specificities and conserved structural features could suggest that there is a pattern of parallel evolution in response to the immediate environment and the lifestyle of the bacteria. We propose an evolutionary path connecting all the AHL and β-lactamacylases based on their oligomeric and activity relationship (Fig. 7). The varied specificity of acylases for different AHLs corroborates very well with the complexity of quorum sensing mechanisms in various bacteria. *Pseudomonas aeruginosa* produces two different AHL signals – C_4_ and 3-oxo-C_12_-HSL, in different ratios while growing in biofilm or planktonic states. In relation, the bacterium exhibits at least four AHL acylases, which might play critical roles in regulating its behaviour (23, 43). The *S. loihica* strain in the present study is the inhabitant of a dynamic microbial mat community characterized byhigh metabolic rates, geochemical gradients and a wide variety of quorum sensing signals (44, 45). It is probable that*Sl*CGH1 and *Sl*CGH2, with their differential AHL specificity, could represent paralogous genes that might be involved in the internal regulation of quorum sensing, as well as quenching of signals from competing bacteria in the environment.

**Fig. 7.**
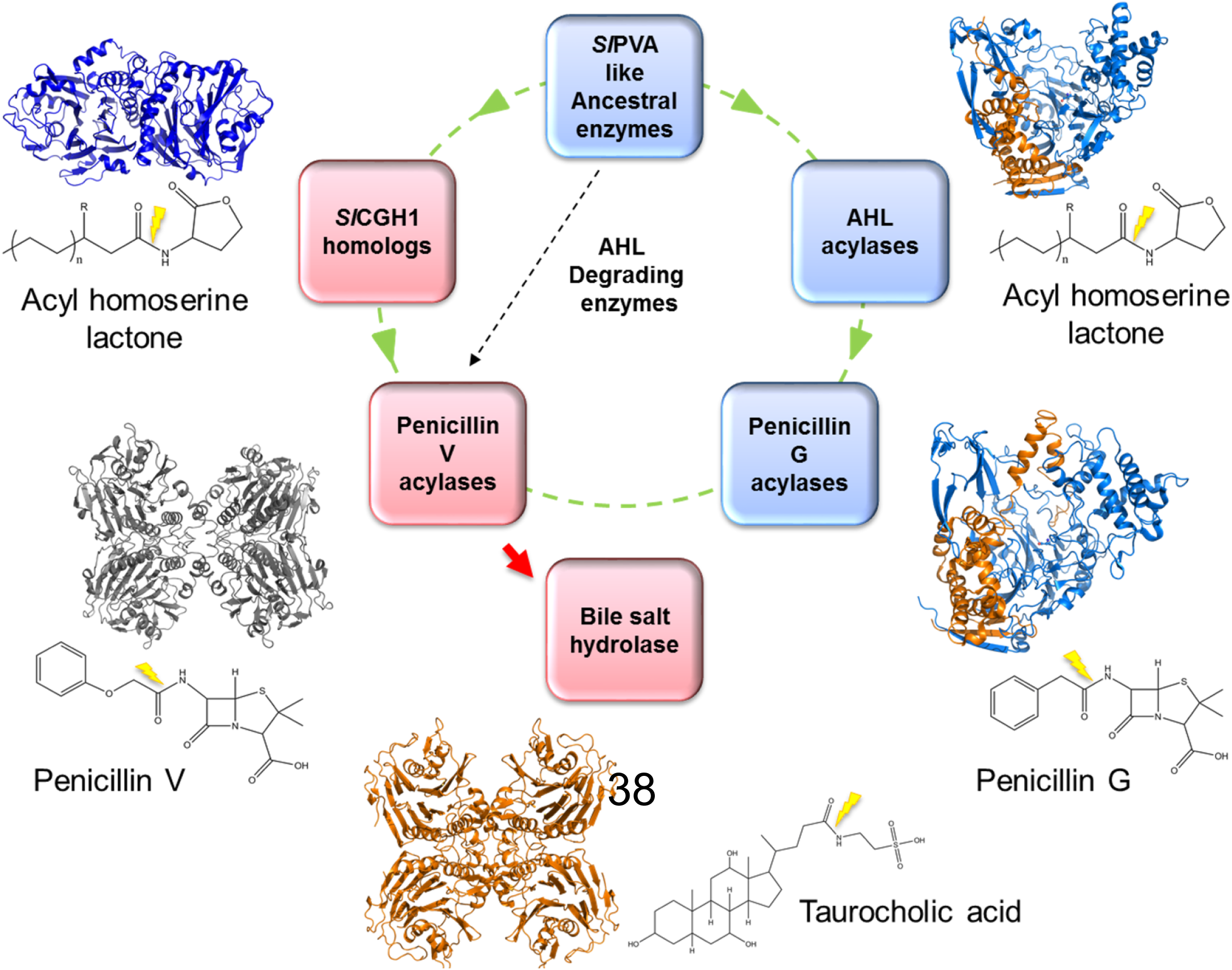
A schematic representation of the probable evolutionary link between different AHL degrading enzyme (green dotted line) (PvdQ, *Kc*PGA, *Pa*PVA and *Sl*CGH1) and β-lactum degrading enzymes (*Kc*PGA, *Pa*PVA, *Sl*PVA), based on substrate specificity and structural overlaps. A promiscious ancestral protein could be similar to heterodimeric *Sl*PVA which shows a wider promiscuity acting on AHLs as well as both aromatic and aliphatic PenVs. Bile salt hydrolase (*Bl*BSH) might be a recently evolved (red arrow) for adaptation in the mammalian gut. Considering the inactivity of *Sl*CGH1 on CGH substrate, independent evolution of PVAs and *Sl*CGH1 homologs remains another possibility (black dotted line).

Many of these acylases that act on AHLs or are evolutionarily related to AHL acylases also exhibit the ability to perform other metabolic functions. PvdQ is an excellent example of dual functionality, being an enzyme of the pyoverdine biosynthesis pathway in *P. aeruginosa*, besides exhibiting AHL acylase activity (46, 47). Many AHL acylases and beta-lactam acylases such as PGA are also suggested to be involved in oligotrophic nutrient scavenging (48, 26). The AHL acylase (MacQ) from *Acidovorax* sp. is able to enhance resistance to multiple penicillins(14). In addition to the bifunctional nature of acylases, recent studies have also brought to light the existence of various novel AHL signals including aryl HSLs (49), branched chain (50) and unsaturated acyl HSLs (51). It is also possible that some acylases might act on yet unidentified endogenous substrates structurally similar to AHLs, to influence and modulate QS-dependent phenotypes (27).There is also a growing popular notion that antibiotics might act as signaling molecules in sub-inhibitory concentrations, controlling gene expression rather than being weapons of destruction (52, 53). The broad-spectrum antibiotic tropodithietic acid (TDA) causes the same regulatory effects in QS comparable to AHLs (54). In combination with the overlapping features of structure and activity between AHL acylases and beta-lactam acylases, this might indicate functional andevolutionarylinks between enzymes involved in signaling and antibiotic resistance.

### Conclusion

Based on the available biochemical and structural information about AHL-active acylases and related enzymes, it appears that they exhibit a complex functional and evolutionary diversity. The present report on cholylglycine hydrolase homologs from the marine *S. loihica* adds to thelist of acylases active on AHLs, while related Ntn hydrolases like BSHs appear to have evolved specificity for other substrates over AHLs. It is reasonable to expect that the discovery of new structurally diverse AHL signals and corresponding quorum quenching enzymes would only add to the complexity. While it might be too ambitious to forge definite links between these enzymes, it can be hoped that future exploration of bacterial QS networks and their regulation could help connect the missinglinks in bacterial communication and thereby better explain the functions of these acylases in relation to bacterial physiology and ecological interactions.

## Methods

A description of methods for protein expression and purification, enzymatic assays including detailed bioluminescence assay, site directed mutagenesis, crystallization of *Sl*CGH1 and ligand complex, docking and molecular dynamics simulation studies, phylogenetic analysis and binding similarity score calculations are provided in SI Materials and Methods.

## Acknowledgement

PDP, VSA and DG thank Council of Scientific and Industrial Research (CSIR), India and YY thanks University Grant Commission (UGC) for the award of Senior Research Fellowship. SR thanks the Department of Science and Technology (DST) for Ramanujan Fellowship and SERB-Young Investigator Grant. All data collections were performed at beam line PXBL21 at RRCAT, BL14 at ESRF, XRD1 at ELETTRA. The coordinates for structure for *Sl*CGH1 native and mutant has been deposited in the PDB under the accession codes **5X8Z** and **5X9I** respectively.

## Supplementary material and method

### Cloning, protein expression and purification of *Sl*CGH1 and *Sl*CGH2

The plasmids, primers and strains used in the study are listed in TableS1. *Sl*CGH1 and *Sl*CGH2 were cloned from the genomic DNA of *S.loihica* into pET22b vector (Invitrogen) between NdeI and XhoI restriction sites, and expressed in *Escherichia coli* BL21 star (DE3) cells with a C-terminal His_6_-tag. Protein expression was induced at OD_600_ = 0.5 by adding 0.2 mM isopropyl-β-D-thiogalactopyranoside (IPTG) to the culture (grown at 37°C for 3-4 h), followed by overnight incubation at 16°C. The harvested cells were resuspended in lysis buffer (25mM Tris-Cl pH 7.0, 300mM NaCl, 10mM MgCl2, 2mM β-mercaptoethanol and 0.1% Triton) and distrupted by sonication. The supernatant was then applied to a HIS-Select Ni^2+^ affinity column and the bound protein eluted with 250 mM imidazole. The protein was further purified to homogeneity using an ENRich^TM^ 650 (BioRad) size exclusion column. SDS-PAGE gel showed a single band of *Sl*CGH1corresponding to 37 KDa molecular mass. Oligomeric state and native molecular weight of *Sl*CGH1 were analyzed using size exclusion Chromatography and MALDI-TOF-MS.

### Screening of enzyme activity

*Sl*CGH1 and *Sl*CGH2 were screened for their activities against standard CGH substrates and selected β-lactam antibiotics. Hydrolysis of penicillin V, penicillin G, amipicillin, carbenicillin and cephalosporin C was assayed by the pDAB method (1, 2). Bile salt hydrolysis (Glycodeoxycholic acid and Taurodeoxycholic acid) was checked using ninhydrin reagent (3). Protease and lipase activities were also screened using methods described by Enyard (4) and Winkler and Stuckmann (5) respectively with slight modifications.

Estimation of AHL degradation was carried out using lux-based bioluminesence assay based on two biosensor strains, *E. coli* pSB401 for C6 to C8-HSLs and *E. coli* pSB1075 for C_10_ to C_12_-HSL (6–8). the protein (20 µg per reaction) was incubated with three different concentrations of each AHL substrates, 15, 25 and 100 µM from a 5 mM stock in DMSO, for 4 h at 30°C in a 96-well plate. The final reaction volume was made up to 100µl using 50 mM sodium phosphate buffer (pH 7.0) with 1mM DTT (6, 7). After incubation, 100 µl of 10 mM PBS (phosphate buffered saline, pH 7.0) was added along with another 100 µl of the 1:100 diluted biosensor culture to appropriate AHL samples and incubated for 6 h at 30°C. Heat-inactivated enzyme was used as control. Bioluminescence was measured in a 96-well white solid plate in a Varioskan^®^ Flash spectral scanning multimode reader driven by SkanIt Software (Thermo Scientific). OD_600_ was also measured in a transparent 96-well plate using the same system, to monitor the bacterial growth. Results were expressed in terms of relative light unit (RLU; LU/OD_600_) obtained from the ratio of the sample readings and the corresponding OD_600_, in relation to the control (control RLU taken as 100%).

### Analysis of AHL degradation by HR-MS spectroscopy

To elucidate and confirm AHL acylase activity, High resolution mass spectroscopy was carried out based on the method described by Huang et al. (9) with slight modifications. Reaction products were analyzed using a C18 ultra-aqueous reverse-phase column fitted to a Q-Exactive system with Accela 1250 Pump and PDA detector from Thermo Scientific. Samples were run for 6 min and the mobile phase consisted of 50 % methanol (methanol-water with 1% acetic acid) for the initial 1 min and linearly increased to 80 % methanol. Samples were diluted in methanol before application. The search criteria for chemical formula determination were limited to a mass error below 5ppm.

### Crystallization and data collection

*Sl*CGH1 was crystallized by vapor diffusion using the hanging drop approach. Diffraction quality crystals were obtained in an optimized condition consisting of 0.2 mM sodium sulphate, 20 % PEG3350 and 5% MPD, with 2 ul protein (20 mg/ml *Sl*CGH1) and 2ul precipitant in the drop. The crystal was soaked in a cryoprotectant solution (30% glycerol) and mounted on a cryo loop, followed by immediate flash-freezing in liquid nitrogen. Diffraction data were collected at Elettra synchrotron facility, Italy. Diffraction data of *Sl*CGH1 crystals were integrated and indexed using XDS (10) and SCALA (11). Co-crystallization of native *Sl*CGH1 and C1S mutant with different AHLs was explored to capture a complex of the enzyme with catalytic reaction intermediates. The protein was pre-incubated in solution with 1-2mM concentration of the substrates for 1 h and kept for crystallization in the same condition as explained above.

### Structure determination

Phase determination was done by molecular replacement using Phaser ver 2.5.6 (12) with the structure of choloylglycine hydrolase from *Bacteroides thetaiotamicron* (PDB code: 3HBC, 22% sequence identity) as the template. Manual model building and refinement were done using COOT and Refmac5 (CCP4 suite). Molecular graphics were made using PyMol (The PyMOL Molecular Graphics System,Version 1.7.4, Schrödinger, LLC).

### Site directed mutagenesis, docking and simulation studies

To ascertain the role of Cys1 as the N-terminal nucleophile, we mutated Cys1 to Ala (C1A) in *SlCGH*1. Another mutation Cys1 to Ser (C1S) was also constructed for comparison with AHL acylases which are Ser Ntn-hydrolases. The mutants were constructed using ExTaq site-directed mutagenesis kit (TaKaRa) and expressed in *E. coli* BL21 cells. Protein expression and purification was performed as detailed for native *Sl*CGH1 above.

Docking analysis was carried out using *Glide* software (Schrodinger Suite 9.0.01, Schrodinger, Inc. Portland USA). 3D conformation of 3-oxo-C_10_-HSL, C_8_-HSL and C_6_-HSL were obtained from PubChem compound database (13) with CID 5282982, CID 6914579, and CID 10058590 respectively and further prepared using the LigPrep. A grid of 26 Å around the nucleophile cysteine was generated using the default parameters. Based on total ligand-receptor interaction energy as Glidescore which includes van der Waals energy and electrostatic energy etc, the best pose was selected.

Molecular Dynamics simulation was carried out using Desmond utility of Maestro, Schrodinger. All the default parameters were used to carry out simulation for 1.2 ns. Chain A of the structure was used as the initial model. In order to explore protein-ligand interaction, three molecular dynamics simulations (MDS); apo-SlCGH1, C1S_3-oxo-octanoic acid and PvdQ_3-oxododecanoic acid (PDB ID: 2wyc) were performed.

### Phylogenetic analysis and Binding similarity score

For phylogenetic analysis (14), protein sequences of 7 structurally characterized CGHs, namely *Bacillus sphaericus* (BspPVA), *B. subtilis* (BsuPVA),*Pectobacterium atrosepticum* (PaPVA), *Bacteroides thetaiotamicron*(BtBSH), *Bifidobacterium longum* (BlBSH) *Clostridium perfringens* (CpBSH) and *Lactobacillus salivaris* (LsBSH) along with *Sl*CGH1 sequence were used as ‘templates’ to retrieve CGH homologs sequences through a BLAST search. The search was filtered through a blast score cut off of 300 and an E-value of 0.008 (0.002 in case of *Sl*CGH1 due to limited number of hits). The cut-off limit was set at 60% redundancy (Jalview software) and sequences lacking N-terminal Cys residue were excluded. After multiple alignment using Clustal X, a phylogenetic tree was constructed in Mega6 (15) using the neighbor-joining method (16) with a bootstrap value of 1000. Further improvement in the graphical representation of the tree was done using Figtree, ver1.42 software (http://tree.bio.ed.ac.uk/software/figtree/).

Panigrahi et al. (14) developed a Binding Site Similarity (BSS) scoring system to estimate quantitatively similarity of each CGH family member based on residue conservation with the binding sites of each of six template CGH enzymes (*Bl*BSH, *Cp*BSH, *Bt*BSH, *Bsp*PVA, *Bsu*PVA and *Pa*PVA). With the addition of *Sl*CGH1 as another template, the score of the query sequence with the *i*th template (*i*=1–7) was calculated as:

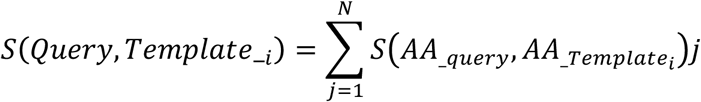

where, *S(AA_query,AA_Tempate_i)j* is the similarity score between amino acid residues of the query and the *i*th template sequence, at the *j*th position of the binding site profile.

## Supplementary Informations

**Fig. S1.**
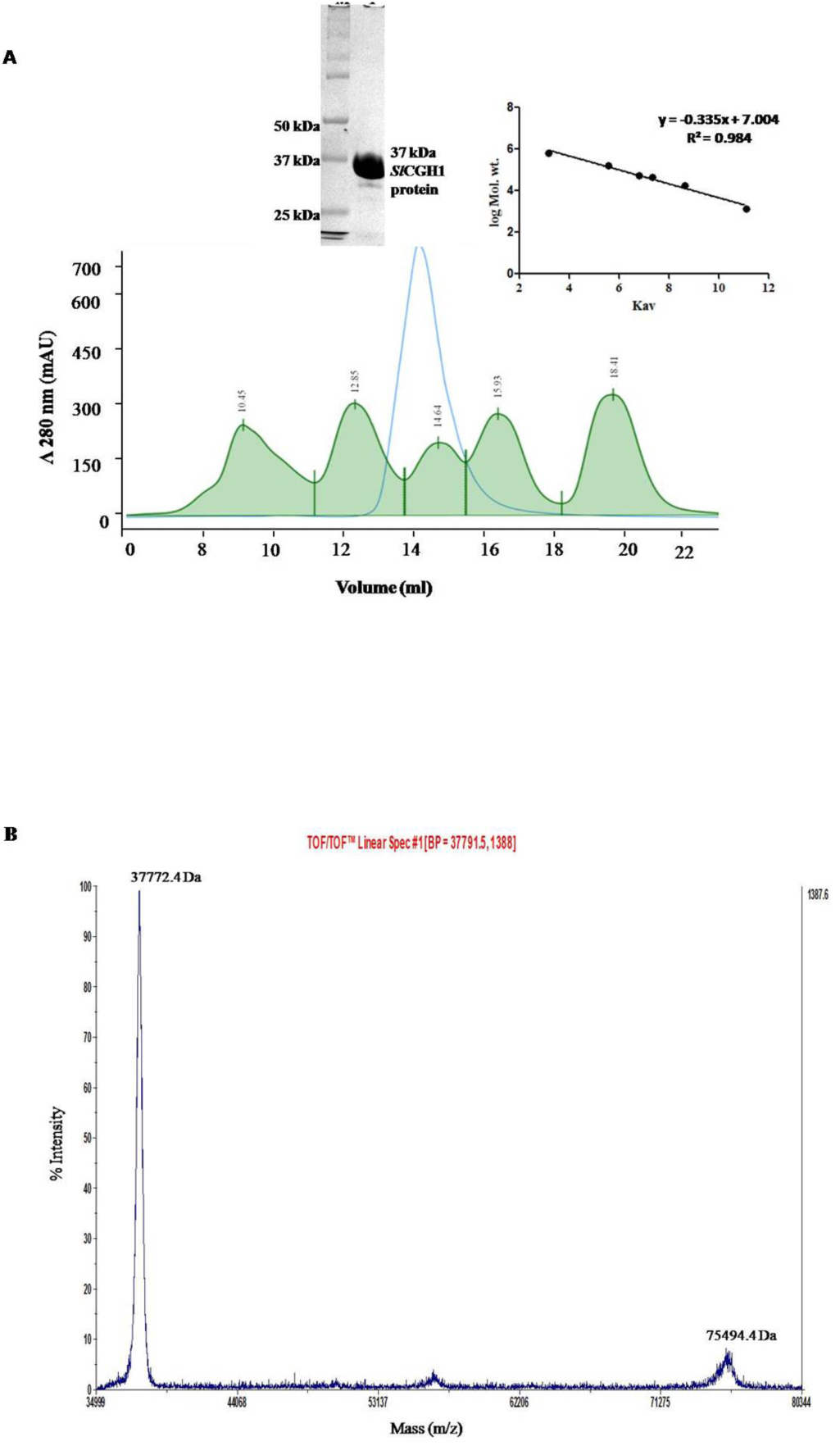
A) Gel filtration chromatography profile of SlCGH1. Purified SlCGH1 of 37 Kda shown against SlCGH1 peak. B) MALDI MS spectrum showing subunit molecular weight of 37 KDa and a dimeric native molecular weight of 75 KDa.

**Fig. S2.**
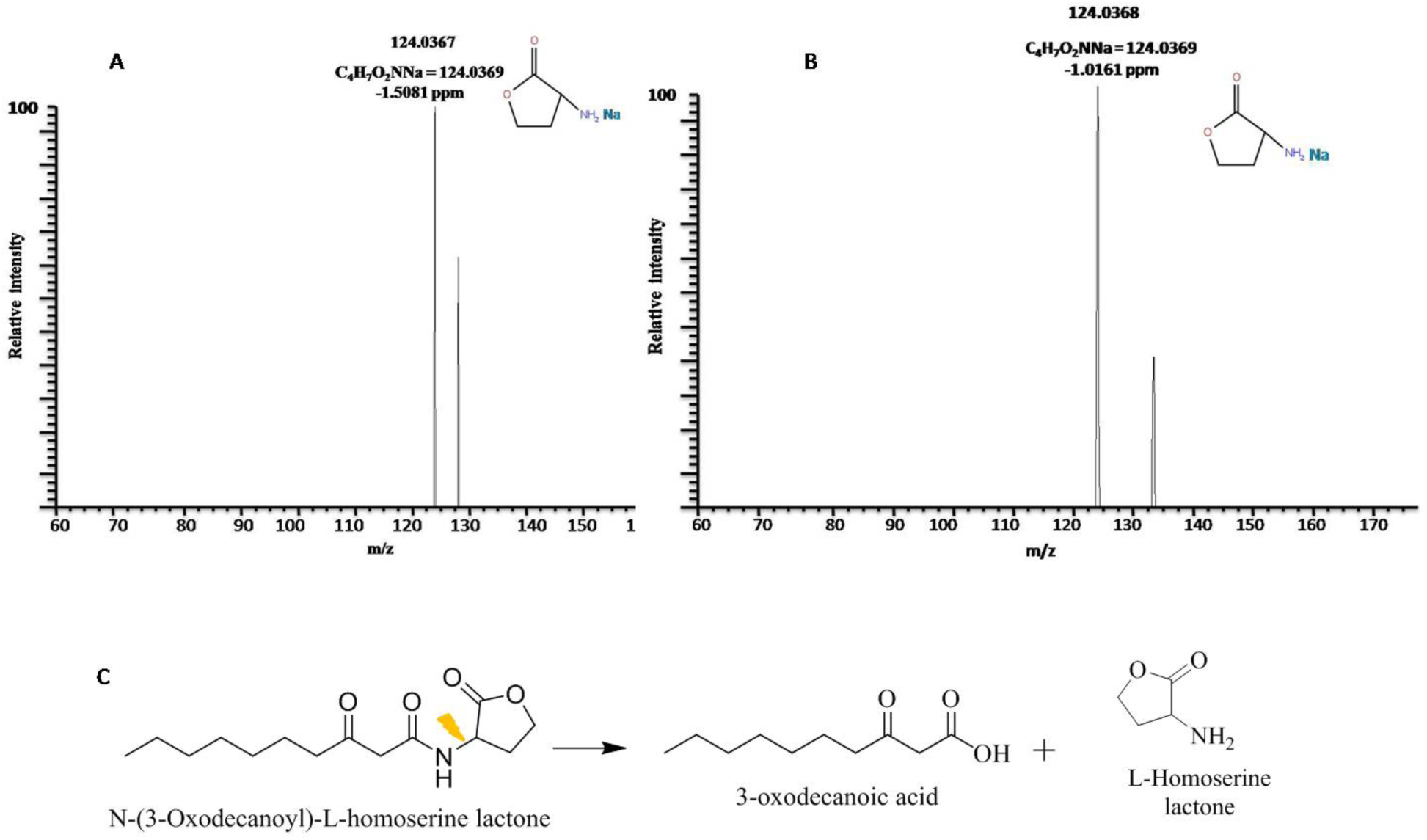
HR-MS spectrometry analysis of A) 3-oxo-C10-HSL degradation by SlCGH1 and B) 3-oxo-C14-HSL degradation by SlCGH2. C) Hydrolysis reaction catalyzed by SlCGH1 (displayed here with 3-oxo-decanoyl HSL as substrate)

**Fig. S3.**
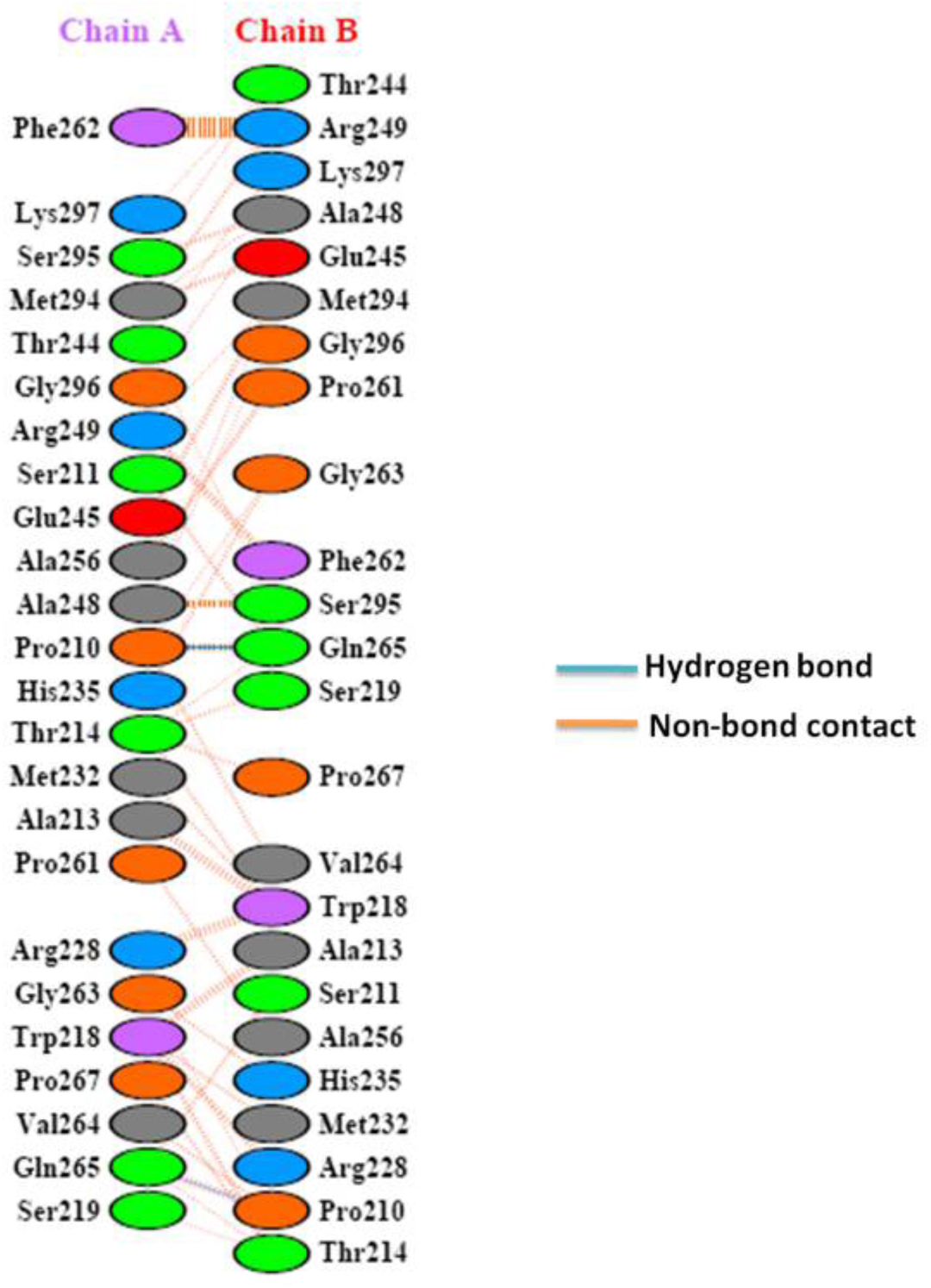
Interface residues interaction between two monomers of a *Sl*CGH1 dimer generated by PDBsum.

**Fig. S4.**
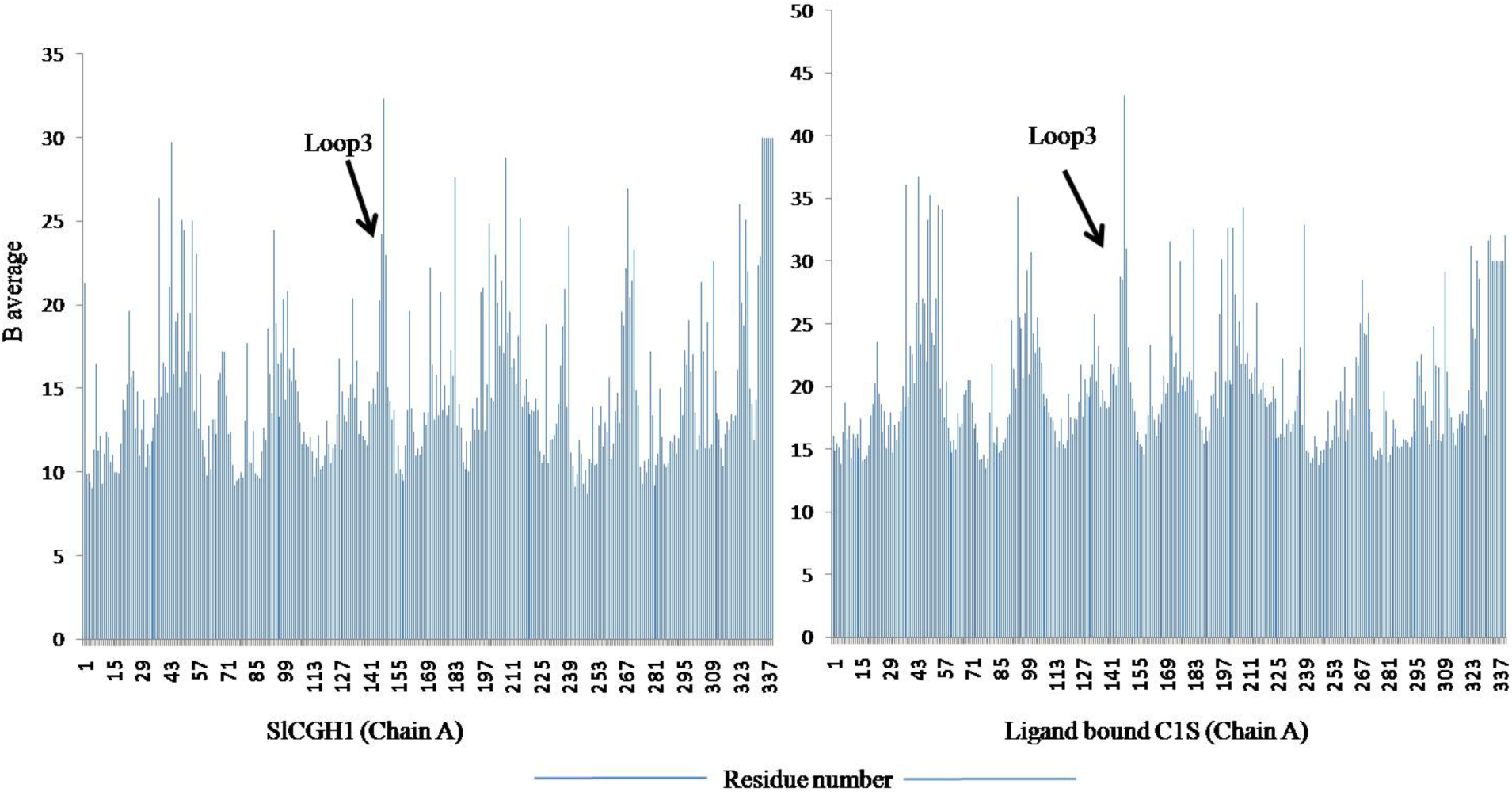
Bav-factor value distribution. A) *Sl*CGH1 B) ligand bound C1S. High B-factor was observed in loop3, centered on residues 147-149, in both apo-*Sl*CGH1 and ligand bound C1S with values increasing from apo to ligand bound structure.

**Fig. S5.**
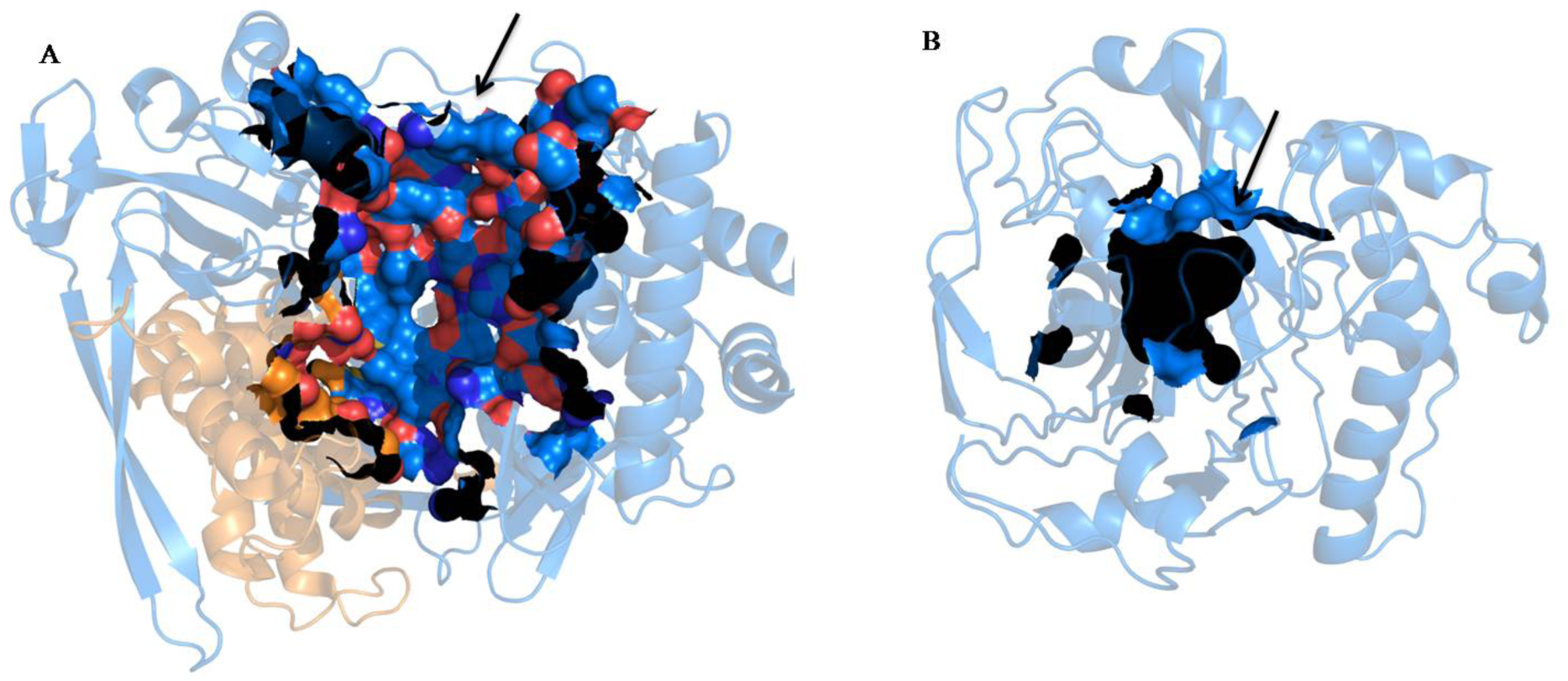
Superimposed structures of apo-*Sl*CGH1 (blue) and ligand bound C1S (red), reveals a low RMSD of 0.083 Å.

**Fig S6.**
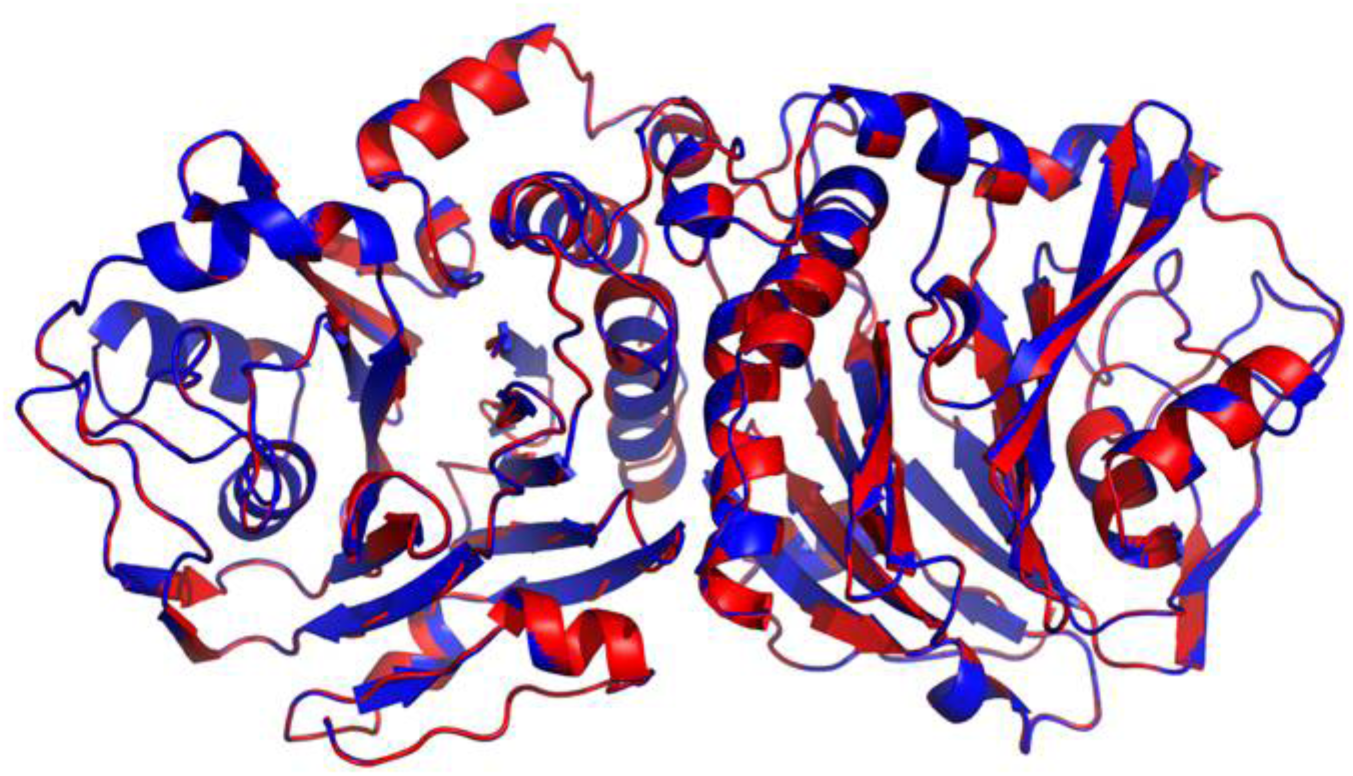
Comparison between active site grooves of SlCGH1 (A) and PvdQ (B, PDB ID 2wyc) using CASTp server (http://sts.bioengr.uic.edu/castp). Active site in PvdQ shows an exposed groove while SlCGH1 exhibits an enclosed active site groove (arrow marked).

**Fig. S7.**
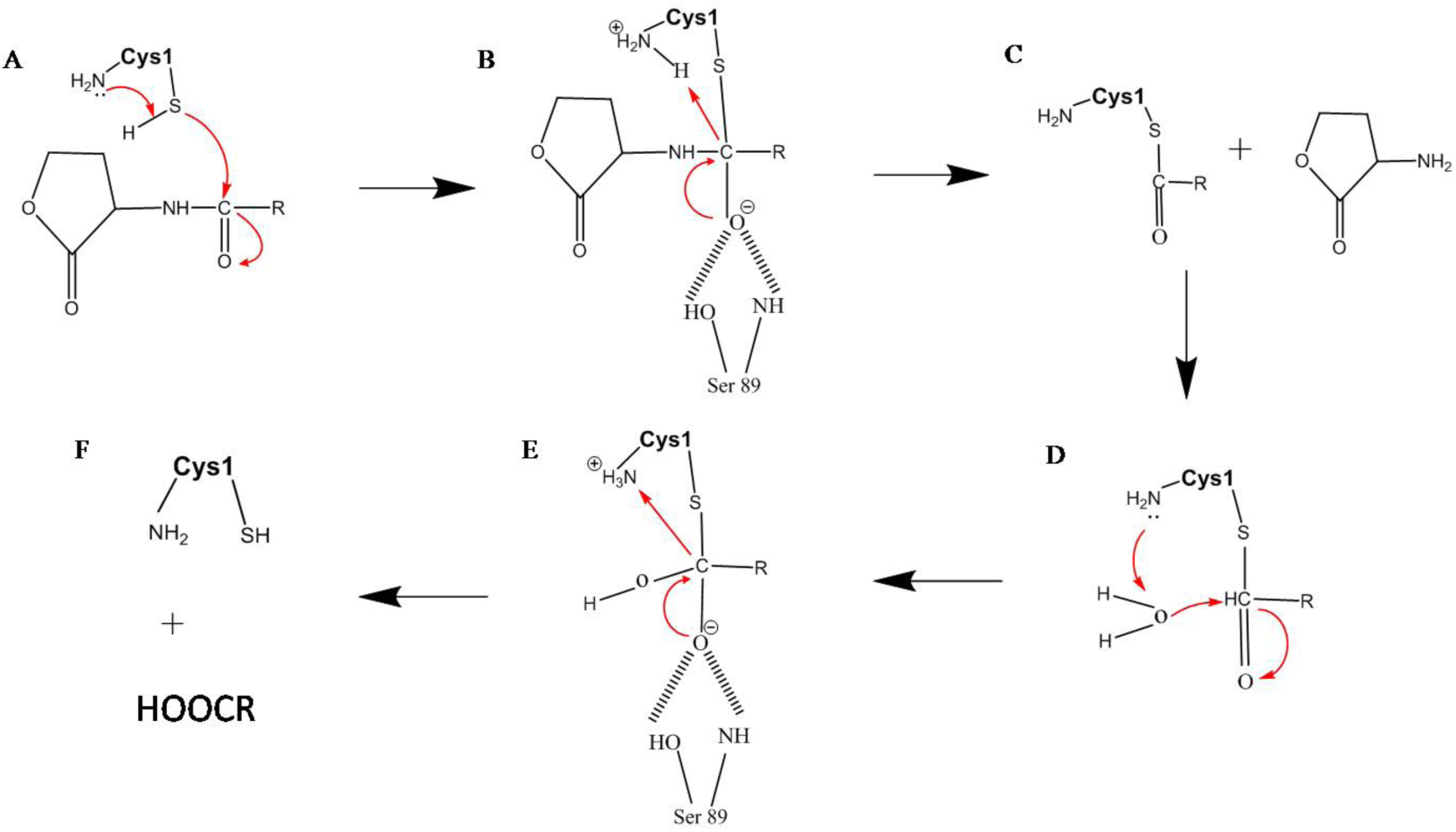
*Sl*CGH1 follows the mechanism of catalysis conserved for Cys-Ntn hydrolases. A) The reaction starts with a direct proton transfer from the Sγ to the a-amino of Cys1 leading to the activation of the nucleophile. B) Sγ makes the nucleophilic attack on the carbonyl carbon of the substrate, resulting to the formation of the tetrahedral intermediate which is stabilized by Ser89 forming the oxyanion hole configuration. C) The α-amino group of Cys donates proton to the nitrogen of the scissile amide bond of the substrate leading to the discharge of the amino part of the substrate. D) The Nα of Cys1 makes Wat1 nucleophilic to make the second nucleophilic attack on the carbonyl carbon of the acyl-enzyme complex. E) A second tetrahedral complex is formed which collapse to release the acyl chain completing the reaction (F).

**Fig S8.**
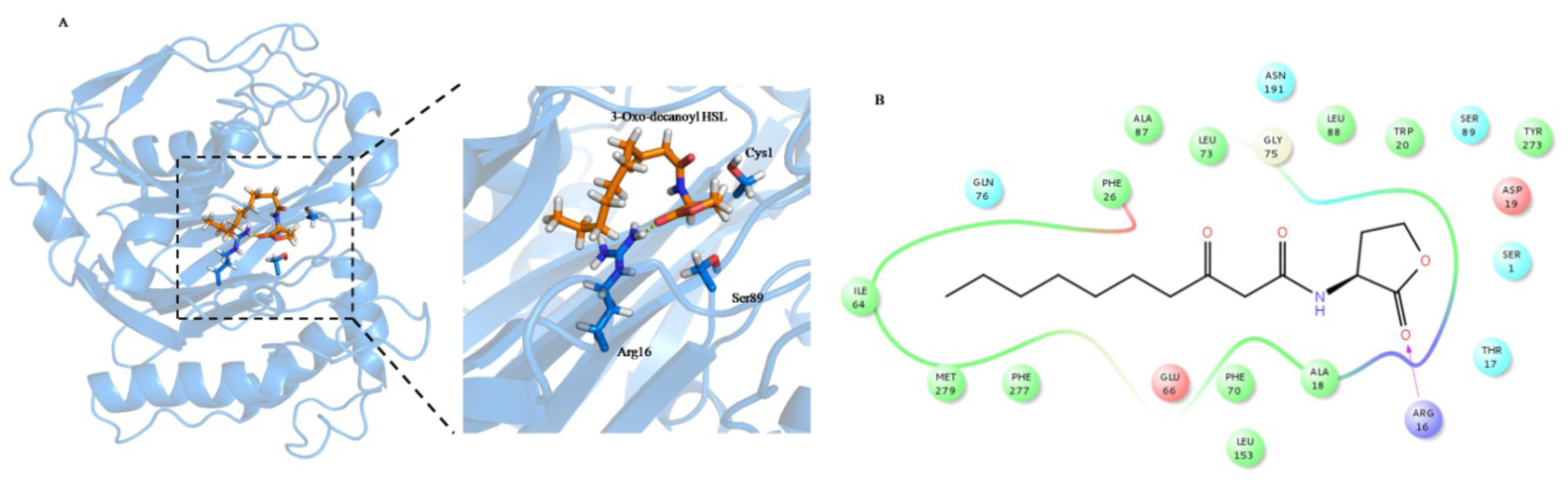
A) Cartoon representation of 3-oxo-C10-HSL docked structure of SlCGH1. 3-Oxo-decanoyl HSL shown as orange and interacting active site residues, Cys1, Arg16 and Ser89. B) 2D representation of A).

**Fig S9.**
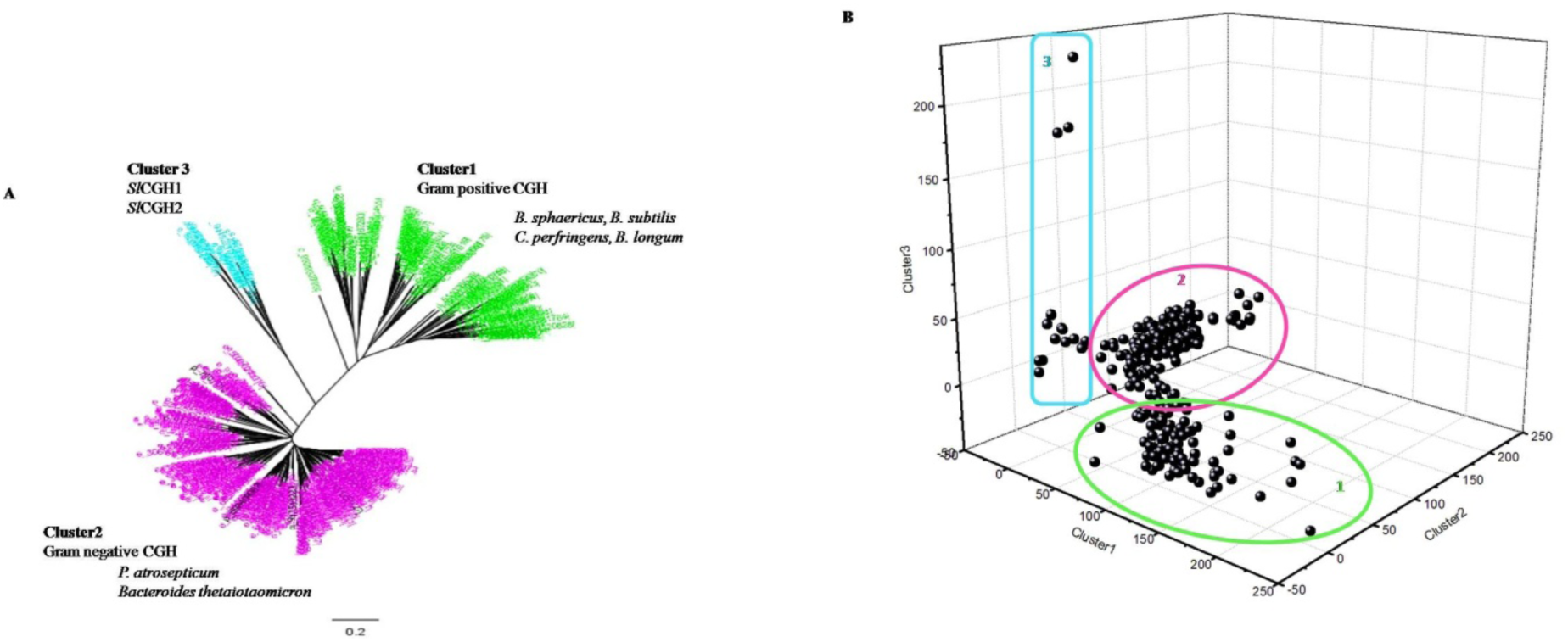
A) Phylogenetic study shows three different well segregated clusters. Cluster1 represents homologs of Gram positive CGHs; Cluster2 Gram negative CGH homologs and Cluster3 SlCGH1 homologs including SlCGH2. Cluster3 is also predominantly marine bacterial CGHs. B) BSS score of CGH homologs presented as 3D graph, showing three clusters of enzymes.

## Tables

**Table S1.** Bacterial strains and plasmids used in the present study

|  | Description | Source |
| --- | --- | --- |
| **Strain** |  |  |
| *Shewanella loihica*-PV4 | CGH host isolated from iron-rich microbial mat of Loihi Seamount in the Pacific Ocean | Gao et al 2006 |
| *E. coli* DH5α | F- 80dlacZ M15 (lacZYA-argF) U169 recA1 endA1 hsdR17(rk-, mk+) phoA supE44 -thi-1 gyrA96 relA1 | Promega |
| *E. coli* BL21 star | F<sup>-</sup>, ompT, hsdSB (rB<sup>-</sup>, mB<sup>-</sup>), dcm, gal, $\lambda$(DE3) | Promega |
| *E. coli* pSB401 | AHL biosensor, Tet<sup>r</sup> | Winson et al 1998 |
| *E. coli* pSB1075 | AHL biosensor, Tet<sup>r</sup> | Winson et al 1998 |
| **Plasmid** |  |  |
| pET22b (+) | Cloning vector; Ap<sup>r</sup> | Novagen |

**Table S2.** Data collection statistics of SlCGH1 crystal and Mutant C1S cocrystalized with 3-oxo-C_8_-HSL (Data in parantheses correspond to the outer shell).

| **X-Ray Diffraction And Data Collection** | ***S/CGH1*** | **Mutant C1S** |
| --- | --- | --- |
| X-ray source | ELETTRA Synchrotron, Italy | ESRF , France |
| Space group | I 12<sub>1</sub> | C12<sub>1</sub> |
| Temperature | 100 K | 100K |
| Resolution range | 48.8-1.8 Å | 50-1.5 Å |
| Unit cell parameters | a=99.4 , b= 52.34, c= 140.07 ;<br>$\alpha = \gamma = 90.00$, $\beta = 103.81$ | a=150.78 , b= 52.42, c= 99.20;<br>$\alpha = \gamma = 90.00$, $\beta = 115.96$ |
| Molecules per asymmetric unit | 2 | 2 |
| Matthews coefficient (Å<sup>3</sup> Da<sup>-1</sup>) | 2.33 | 2.38 |
| Solvent content (%) | 48.58 | 48.39 |
| Total no. of reflections | 386710 (18943) | 177994(15294) |
| No. of unique reflections | 109332 (5258) | 36355(3127) |
| Multiplicity | 3.5 (3.6) | 4.9(4.9) |
| Completeness (%) | 97.6 | 100 |
| Average I/$\sigma$(I) | 13.8 (3.3) | 12.2(5.1) |
| R-merge (all I+ and I-) | 0.060 (0.0309) | 0.084(0.276) |

**Table S3.** Summary of data collection and refinement statistics

| **Crystal system / Space group** | ***SICGH1*** | **Mutant C1S** |
| --- | --- | --- |
| Resolution range (Å) | 1.8 Å | 1.5 Å |
| Completeness (%) | 97.5 | 100 |
| Number of unique reflections | 109332 | 36355 |
| Rsym (%) | 6.0 | 8.4 |
| **<u>Refinement statistics</u>** |  |  |
| R<sub>cryst</sub> | 0.190 | 0.185 |
| R<sub>free</sub> | 0.216 | 0.208 |
| Number of protein atoms | 5322 | 5445 |
| Number of water molecules | 172 | 175 |
| Overall B value (Å<sup>2</sup>) | 15.487 | 20.656 |
| **<u>Root mean square deviation</u>** |  |  |
| Bond length (Å) | 0.0253 | 0.219 |
| Bond angle (°) | 2.2400 | 2.0757 |
| Dihedral angles | 0.152 | 0.135 |
| **<u>Ramachandran plot (% residues)</u>** |  |  |
| Most favoured region | 96.4 % | 96.3 % |
| Allowed region | 3.1 % | 3.7 % |
| Outlier region | 0.4 % | 0 |

**Table S4.** Structural alignment of SlCGH1 with other Ntn hydrolases using DALI server (Holm and Rosenstrom 2010).

| PDB ID | Z-score | RMSD | L-Ali | NRES | % Identity | Protein |
| --- | --- | --- | --- | --- | --- | --- |
| 3HBC | 33.8 | 2.4 | 295 | 309 | 20 | CGH (*B. thetaiotamicron*) |
| 4WL2 | 33.8 | 2.6 | 309 | 346 | 18 | PVA (*P. atrosepticum*) |
| 2PVA | 31.4 | 2.6 | 293 | 332 | 18 | PVA (*B. sphaericus*) |
| 2OQC | 31.1 | 2.7 | 293 | 317 | 19 | PVA (*B. subtilis*) |
| 2BJF | 31.1 | 2.7 | 291 | 328 | 16 | BSH (*C.perfingens*) |
| 2HEZ | 28.9 | 2.8 | 283 | 316 | 14 | BSH (*B. longum*) |
| 3FGT | 16 | 3.3 | 231 | 344 | 7 | Phospholipase B-like |
| 1GK1 | 15.3 | 3.4 | 224 | 522 | 6 | Cephalosporin acylase |
| 4M1J | 14.3 | 3.7 | 227 | 547 | 7 | AHL acylasePvdQ |
| 3M1O | 13.9 | 3.7 | 224 | 551 | 10 | Penicillin G acylaseα subunit |

**Table S5.** Properties of different AHL substrates and results of docking with *Sl*CGH1 structure (AlogP = hydrophobicity, SA = surface area, Nadist = Nucleophilic attack distance between SH group of cys1 and carbonyl carbon atom of AHL).

| Ligand<br>(AHL) | MW | PolarSA | AlogP | Rotatable<br>bo<br>nds | H-bond<br>don<br>ors | H-bond<br>accept<br>ors | Glidescore | Nadist(<br>Å) |
| --- | --- | --- | --- | --- | --- | --- | --- | --- |
| C<sub>6</sub><br>HSL | 199.3 | 55.4 | 1.133 | 6 | 1 | 3 | -2.84 | 8.65 |
| C<sub>8</sub><br>HSL | 227.3 | 55.4 | 2.6 | 7 | 1 | 3 | -2.81 | 6.62 |
| 3-oxo-<br>C<sub>10</sub><br>HSL | 255.4 | 55.4 | 2.958 | 10 | 1 | 3 | -4.24 | 5.9 |

## References

1. Roodveldt C, Tawfik DS (2005) Shared promiscuous activities and evolutionary features in various members of the amidohydrolase superfamily. Biochemistry 44(38):12728–12736.

2. Park SY, Kang HO, Jang HS, Lee JK, Koo BT, Yum DY (2005) Identification of extracellular N-acylhomoserine lactone acylase from a *Streptomyces* sp. and its application to quorum quenching. Appl. Environ. Microbiol. 71:2632–2641.

3. Krzeslak J, Wahjudi M & Quax WJ. (2007). Quorum quenching acylases in Pseudomonasaeruginosa.InPseudomonas: a Model System in Biology, vol. 5, pp. 429–449. Edited by J.-L. Ramos & A. Filloux. Netherlands:

4. Mukherji R, Varshney NK, Panigrahi P, Suresh CG, Prabhune A (2014) A new role for penicillin acylases: Degradation of acyl homoserine lactone quorum sensing signals by *Kluyveracitrophila* penicillin G acylase. Enzyme Microb Technol 56:1–7.

5. Avinash VS, Utari PD, Ramasamy S, Merkerk VR, Quax W, Pundle A(2017) Penicillin V acylases from gram-negative bacteria degrade N-acylhomoserine lactones and attenuate virulence in *Pseudomonas aeruginosa*. Appl Microbiol Biotechnol 101:2383–2395.

6. Avinash VS, Pundle AV, Ramasamy S, Suresh CG (2016) Penicillin acylases revisited: importance beyond their industrial utility. Crit Rev Biotechnol 36:303–316.

7. Miller MB, Bassler BL (2001) Ensing in. Annu Rev Microbiol 55:165–99.

8. Whitehead N a, Barnard a M, Slater H, Simpson NJ, Salmond GP (2001) Quorum-sensing in Gram-negative bacteria. FEMS Microbiol Rev 25:365–404.

9. Uroz S, Dessaux Y, Oger P (2009) Quorum sensing and quorum quenching: the yin and yang of bacterial communication. Chembiochem 10:205–16.

10. Fetzner S (2014) Quorum quenching enzymes. J Biotechnol 201:2–14.

11. Grandclément C, Tannières M, Moréra S, Dessaux Y, Faure D (2015) Quorum quenching: Role in nature and applied developments. FEMS Microbiol Rev 40:86–116.

12. Elias M, Tawfik DS (2012) Divergence and convergence in enzyme evolution: Parallel evolution of paraoxonases from quorum-quenching lactonases. J Biol Chem 287:11–20.

13. Bokhove M, Jimenez PN, Quax WJ, Dijkstra BW (2010) The quorum-quenching N-acyl homoserine lactone acylase PvdQ is an Ntn-hydrolase with an unusual substrate-binding pocket. Proc Natl Acad Sci 107:686–691.

14. Kusada H, Tamaki H, Kamagata Y, Hanada S, Kimura N (2017) A novel quorum-quenching enzyme mediates antibiotic resistance. Appl Environ Microbiol (April):AEM.00080-17.

15. Suresh CG, Pundle AV, SivaRaman H, Rao KN, Brannigan JA, McVey CE, Verma CS, Dauter Z, Dodson EJ, Dodson GG (1999) Penicillin V acylase crystal structure reveals new Ntn-hydrolase family members. Nat Struct Biol 6:414–416.

16. Gao H, Obraztova A, Stewart N, Popa R, Fredrickson KJ, Tiedje MJ, Nealson HK, Zhou J (2006) *Shewanella loihica* sp. nov., isolated from iron-rich microbial mats in the Pacific Ocean. Int J Syst Evol Microbiol 56:1911–1916.

17. Avinash VS (2015) Penicillin V Acylases From Gram-Negative Bacteria : Biochemical And Structural Aspects. Int J Biol Macromol 79:1–7.

18. Panigrahi P, Sule M, Sharma R, Ramasamy S, Suresh CG (2014) An improved method for specificity annotation shows a distinct evolutionary divergence among the microbial enzymes of the cholylglycine hydrolase family. Microbiol 160:1162–1174.

19. Lambert JM, Bongers RS, De Vos WM, Kleerebezem M (2008) Functional analysis of four bile salt hydrolase and penicillin acylase family members in *Lactobacillus plantarum* WCFS1. Appl Environ Microbiol 74:4719–4726.

20. Avinash VS, Vellore Sunder, Priyabrata P, Chand D, Pundle A, Suresh CG. (2016) Structural analysis of a penicillin V acylase from *Pectobacterium atrosepticum* confirms the importance of two Trp residues for activity and specificity. J Struct Biol 193:85–94.

21. Morohoshi T, Nakazawa S, Ebata A, Kato N, Ikeda T (2008) Identification and characterization of N-acylhomoserine lactone-acylase from the fish intestinal *Shewanella* sp. strain MIB015. Biosci Biotechnol Biochem 72:1887–93.

22. Huang JJ, Han JI, Zhang LH, Leadbetter JR (2003) Utilization of acyl-homoserine lactone quorum signals for growth by a soil pseudomonad and *Pseudomonas aeruginosa* PAO1. Appl Environ Microbiol 69:5941–5949.

23. Huang JJ, Petersen A, Whiteley M, Leadbetter JR (2006) Identification of QuiP, the product of gene PA as the second acyl-homoserine lactone acylase of *Pseudomonasaeruginosa* PAO1. Appl Environ Microbiol 72:1190–1197.

24. Romero M, Diggle SP, Heeb S, Camara M, Otero A (2008) Quorum quenching activity in Anabaena sp. PCC 7120: identification of AiiC, a novel AHL-acylase. FEMS Microbiol Lett 280:73–80.

25. Koch G, Nadal-Jimenez P, Cool RH, Quax WJ (2014) *Deinococcusradiodurans* can interfere with quorum sensing by producing an AHL-acylase and an AHL-lactonase. FEMS Microbiol Lett 356:62–70.

26. Lin YH, Xu JL, Hu J, Wang LH, Ong SL, Leadbetter JR, Zhang LH (2003) Acyl-homoserine lactone acylase from *Ralstonia* strain XJ12B represents a novel and potent class of quorum-quenching enzymes. Mol Microbiol 47:849–860.

27. Shepherd RW, Lindow SE (2009) Two Dissimilar N-Acyl-Homoserine Lactone Acylases of *Pseudomonassyringae* Influence Colony and Biofilm Morphology. Appl Environ Microbiol 75:45–53.

28. Kumar RS, Brannigan AJ, Prabhune AA, Pundle VA, Dodson GG, Dodson E, Suresh CG (2006) Structural and functional analysis of a conjugated bile salt hydrolase from *Bifidobacterium longum* reveals an evolutionary relationship with penicillin V acylase. J Biol Chem 281:32516–32525.

29. Stellwag EJ, Hylemon PB (1976) Purification and characterization of bile-salt hydrolase from *Bacteroidesfragilis subsp fragilis*. Biochim Biophys Acta 452:165–176.

30. Rossocha M, Schultz-heienbrok R, Moeller H Von, Coleman JP, Saenger W (2005) Conjugated bile acid hydrolase is a tetrameric N-terminal thiol hydrolase with specific recognition of its cholyl but not of its tauryl product. Biochem 2:5739–5748.

31. Palermo G, Bauer I, Campomanes P, Cavalli A, Armirotti A, Girotto S, Rothlisberger U and De Vivo M (2015) Keys to Lipid Selection in Fatty Acid Amide Hydrolase Catalysis: Structural Flexibility, Gating Residues and Multiple Binding Pockets. PLOS Comput Biol 11:e1004231.

32. Duggleby HJ, Tolley SP, Hill CP, Dodson EJ, Dodson G, Moody PCE (1995) Penicillin acylase has a single amino acid catalytic center Nature 373:264–268.

33. Oinonen C, Rouvinen J (2000) Structural comparison of Ntn-hydrolases. Prot Sci 9:2329– 2337.

34. Kyeong Sook Choi, Jong Ahn Kim, Hyen Sam Kang (1992) Effects of site-directed mutations on processing and activities of penicillin G acylase from Escherichia coli ATCC 11105. J Bacteriol 174:6270–6276.

35. Noren C, Wang J, Perler F (2000) Dissecting the chemistry of protein splicing and its applications. Angew Chem Int Ed Engl 39:450–466.

36. Lodola A, Branduardi D, Capoferri MVL, Mor M, Piomelli D, Cavalli A (2012) A catalytic mechanism for cysteine N-terminal nucleophile hydrolases, as revealed by free energy simulations. PLoS One 7: e32397.

37. Chand D, Ramasamy S, Suresh CG (2016) A highly active bile salt hydrolase from *Enterococcusfaecalis* shows positive cooperative kinetics. Proc Biochem 2: 263–269.

38. Rossmann M (2008) PhD thesis. Structural analysis of proteins of sphingolipid metabolism, Freie Universitat Berlin, Berlin.

39. Swearingen MC, Sabag-Daigle A, Ahmer BMM (2013) Are there acyl-homoserine lactones within mammalian intestines? J Bacteriol 195:173–179.

40. Ochiai S, Morohoshi T, Kurabeishi A, Shinozaki M, Fujita H, Sawada I, Ikeda T (2013) Production and degradation of N -acylhomoserine 006Cactone quorum Sensing Signal molecules in bacteria isolated from activated sludge. Biosci Biotechnol Biochem 77:2436– 2440.

41. Koch G, Nadal-Jimenez P, Reis CR, Muntendam R, Bokhove M, Melillo E, Dijkstra BW, Cool RH, Quax WJ (2014) Reducing virulence of the human pathogen *Burkholderia* by altering the substrate specificity of the quorum-quenching acylase PvdQ. Proc Nat Acad Sci 111:1568–1573.

42. Czajkowski R, Krzyzanowska D, Karczewska Joanna, Atkinson S, Przysowa J, Lojkowska E, Williams P, Jafra S (2011) Inactivation of AHLs by Ochrobactrum sp. A44 depends on the activity of a novel class of AHL acylase. Environ Microbiol Rep 3:59–68.

43. Krzeslak J, Wahjudi M, Quax WJ (2007) Quorum quenching acylases in Pseudomonas aeruginosa. In: Ramos JL, Filloux A (eds) Pseudomonas. Springer, New York, pp. 429–449.

44. Decho AW, Decho AW, Visscher PT, Ferry JK, Tomohiro H, Lijian P, Kristen MN, Reid RS, Pamela R. (2009) Autoinducers extracted from microbial mats reveal a surprising diversity of N-acylhomoserine lactones (AHLs) and abundance changes that may relate to diel pH. Environ Microbiol 11:409–420.

45. Decho AW, Norman RS, Visscher PT (2010) Quorum sensing in natural environments: emerging views from microbial mats. Trends Microbiol 18:73–80.

46. Ochsner UA, Wilderman PJ, Vasil AI, Vasil ML (2002) GeneChip® expression analysis of the iron starvation response in *Pseudomonas aeruginosa*: Identification of novel pyoverdine biosynthesis genes. Mol Microbiol 45:1277–1287.

47. Lamont IL, Martin LW (2003) Identification and characterization of novel pyoverdine synthesis genes in *Pseudomonas aeruginosa*. Microbiology 149:833–842.

48. Valle F, Balbas P, Merino E, Bolivar F (1991) The role of penicillin amidases in nature and in industry. Trends Biochem Sci 16:36–40.

49. Ahlgren NA, Harwood CS, Schaefer AL, Giraud E, Greenberg EP (2011) Aryl-homoserine lactone quorum sensing in stem-nodulating photosynthetic *Bradyrhizobia*. Proc Natl Acad Sci 108:7183–7188.

50. Lindemanna A, Pessi G, Schaefera L A, Mattmannc EM, Christensend HQuin, Kesslerb A, Henneckeb H, Blackwellc EH, Greenberga EP, and Harwood SC (2011) Isovaleryl-homoserine lactone, an unusual branched-chain quorum-sensing signal from the soybean symbiont *BradyrhizobiumJaponicum*. Proc Nat Acad Sci 108:16765–16770.

51. Shen Q, Gao J, Liu J, Liu S, Liu Z, Wang Y, Guo B, Zhuang X, Zhuang G (2016) A New Acyl-homoserine Lactone Molecule Generated by *Nitrobacterwinogradskyi*. Sci Rep 6:22903.

52. Romero D, Traxler MF, López D, Kolter R (2011) Antibiotics as signal molecules. Chem Rev 111:5492–5505.

53. Struss AK, Pasini P, Flomenhoft D, Shashidhar H, Daunert S (2012) Investigating the effect of antibiotics on quorum sensing with whole-cell biosensing systems. Anal Bioanal Chem 402:3227–3236.

54. Beyersmann PG, Tomasch J, Son K, Stocker R, Göker M, Wagner-Döbler I, Simon M, Brinkhoff T (2017) Dual function of tropodithietic acid as antibiotic and signaling molecule in global gene regulation of the probiotic bacterium *Phaeobacterinhibens*. Sci Rep 7:730.

55. Ashkenazy H, Abadi S, Martz E, Chay O, Mayrose I and Pupko T (2016) ConSurf 2016: an improved methodology to estimate and visualize evolutionary conservation in macromolecules. Nucleic Acids Res 44(W1):W344–W350.

## References

1. Bomstein J, Evans WG (1965) Automated colorimetric determination of 6-amino penicillanic acid in fermentation media. Anal Chem 37:576–578.

2. Shewale JG, Kumar KK, Ambekar GR (1987) Evaluation of determination of 6-amino penicillanic acid by p-dimethylaminobenzaldehyde. Biotechnol Tech 1:69–72.

3. Kumar RS, Brannigan AJ, Prabhune AA, Pundle VA, Dodson GG, Dodson E, Suresh CG (2006) Structural and functional analysis of a conjugated bile salt hydrolase from *Bifidobacterium longum* reveals an evolutionary relationship with penicillin V acylase. J Biol Chem 281:32516–32525.

4. Cupp-Enyard C (2008) Sigma’s Non-specific Protease Activity Assay - Casein as a Substrate. J Vis Exp e899:1–3.

5. Winkler UK, Stuckmann M (1979) Glycogen, hyaluronate, and some other polysaccharides greatly enhance the formation of exolipase by *Serratia marcescens*. J Bacteriol 138:663–670.

6. Steindler L, Venturi V (2007) Detection of quorum-sensing N-acyl homoserine lactone signal molecules by bacterial biosensors. FEMS Microbiol Lett 266:1–9.

7. Winson MK, Swift S, Fish L, Throup PJ, Frieda J, Chhabra, Siri Ram, Bycroft BW, Williams P, Stewart G SAB (1998) Construction and analysis of luxCDABE -based plasmid sensors for investigating N -acyl homoserine lactone-mediated quorum sensing. FEMS Microbiol Lett 163(2):185–192.

8. Wahjudi M, Papaioannou E, Hendrawati O, van Assen AHG, van Merkerk R, Cool RH, Poelarends GJ, Quax WJ (2011) PA0305 of *Pseudomonas aeruginosa* is a quorum quenching acylhomoserine lactone acylase belonging to the Ntn hydrolase superfamily. Microbiol 157:2042–55.

9. Huang JJ, Han JI, Zhang LH, Leadbetter JR (2003) Utilization of acyl-homoserine lactone quorum signals for growth by a soil pseudomonad and *Pseudomonas aeruginosa* PAO1. Appl Environ Microbiol 69:5941–5949.

10. Kabsch W (2010) XDS. Acta CrystallogrD 66:125–132.

11. Evans P (2006) Scaling and assessment of data quality. Acta Crystallogr D 62:72– 82002E.

12. McCoy AJ, Grosse-Kunstleve RW, Adams PD, Winn MD, Storoni LC, Read RJ (2007) Phaser crystallographic software. J Appl Crystallogr 40:658–674

13. Bolton E, Wang Y, Thiessen PA, Bryant SH (2008) PubChem: Integrated platform of small molecules and biological activities, in: Annual Reports in Computational Chemistry, Elsevier, Washington DC, 217–241.

14. Panigrahi P, Sule M, Sharma R, Ramasamy S, Suresh CG (2014) An improved method for specificity annotation shows a distinct evolutionary divergence among the microbial enzymes of the cholylglycine hydrolase family. Microbiol 160:1162–1174.

15. Tamura K, Stecher G, Peterson D, Filipski A, Kumar S (2013) MEGA6: Molecular Evolutionary Genetics Analysis Version 6.0. Mol Biol Evol 30:2725–2729.

16. Saitou N, Nei M (1987) The neighbor-joining method: a new method for reconstructing phylogenetic trees. Mol Biol Evol 4:406–425.

